# Exploration of Novel Biomarkers through a Precision Medicine Approach Using Multi-omics and Brain Organoids in Patients with Atypical Depression and Psychotic Symptoms

**DOI:** 10.1101/2025.04.21.649748

**Authors:** Insook Ahn, Soyeon Chang, Seok-Ho Choi, Jinju Han, Yangsik Kim

## Abstract

Major depressive disorder (MDD) with atypical features accompanied by psychotic symptoms represents a severe and under-researched subtype of depression and severe mental illness, characterized by significant personal and social impact. This study aims to explore novel biomarkers through a precision medicine approach by integrating clinical data, white blood cell (WBC) single-cell RNA sequencing (scRNA-seq), plasma proteomics, and brain organoid models to uncover immunological and neurological alterations in patients with this condition. Patients exhibited elevated stress, anxiety, and depression levels, with increased WBC counts. Plasma proteomic profiling identified an upregulation of proteins implicated in synaptic formation, including Doublecortin-Like Kinase 3 (DCLK3) and Calcyon (CALY), as well as immune-related proteins such as Complement Component 5 (C5). WBC scRNA-seq revealed significant neutrophil and monocyte transcriptomic alterations, suggesting increased inflammation and immune dysregulation. Patient-derived brain organoids display reduced growth and distinct gene expression patterns compared to controls, particularly under dexamethasone-induced stress conditions. Integrating multi-omics data and brain organoid models offers a novel framework for understanding the pathophysiology of psychiatric disorder, which is one of the most complex disorders.

## Introduction

The Global Burden of Disease 2019 showed that mental disorders affected 907.1 million people and accounted for 125.3 million disability-adjusted life years (DALYs), ranking it as the seventh highest contributor to the global disease burden^1^. Mood disorders, following anxiety disorders, affected 279.6 million people (28.8%), with a higher prevalence among women, and accounted for the highest proportion of DALYs (37.3%). Major depressive disorder (MDD), which is one of the most common mood disorders, is classified as treatment-resistant depression in about one-third of cases and causes significant personal distress and social burden^2^. Despite the high prevalence and social impact of mood disorders, there is a lack of objective biomarkers for their diagnosis and treatment, highlighting the need for further research^3^.

Immunological abnormalities have been consistently reported in mood disorders, particularly MDD^3–11^. MDD with immune dysfunction is often less responsive to conventional antidepressant medication use, and patients with distinct immunological abnormalities have shown improvement with the use of infliximab, an anti-TNF immunosuppressant^12^. It is also well established that depressive symptoms can be induced by the pro-inflammatory cytokine such as IFN-^13^. Among the MDD subtypes, patients with atypical and melancholic features reportedly exhibit immunological abnormalities^6–9,14–16^. These specific subtypes may have different immune activation profiles (such as Th1 and Th2 responses), which could necessitate tailored treatment strategies^17,18^. However, clinical studies are yet to show significant differences in drug efficacy based on subtypes^19,20^.

Immunological abnormalities have also been reported in psychotic disorders including schizophrenia, which shares biological abnormalities with mood disorders^21–24^. Common cytokine alterations have been observed in mood disorders and schizophrenia^25,26^, and immune dysfunction has also been reported in studies of MDD accompanied by psychotic symptoms^27,28^. Mood and psychotic symptoms in adulthood are commonly associated with childhood stress, adversity, and inflammatory states^29–32^. Additionally, psychotic experiences, including idea of reference (IOR), are observed in approximately 8% of the general population^33,34^; psychotic symptoms occur in 9.8%– 10.92% of patients with MDD and are associated with more severe depression, higher anxiety levels, and increased suicide risk^35–38^. Thus, we focused on patients with severe mental illness (SMI), particularly those diagnosed with MDD with atypical features accompanied by psychotic symptoms, because this condition is associated with a more severe disease trajectory.

One of the greatest challenges in psychiatric research is the inability to study a living and functional brain that is intact and connected. Psychiatric disorders are triggered by the complex interplay of individual vulnerabilities and environmental stress, including lifetime adversity. These vulnerabilities encompass not only genetic predispositions but also acquired factors, such as stress-induced changes, particularly recently reported brain somatic mutations^39,40^. Therefore, an approach that reflects both a functioning brain and an individual’s biological characteristics is required, and these limitations, arising from a lack of representative models, could potentially be addressed using brain organoids^41–43^. By isolating blood monocytes and using induced pluripotent stem cells (iPSCs) to differentiate and form brain organoids, an in vitro functional brain model can reflect patient genetic and biological characteristics. Additionally, the administration of dexamethasone (DMX), a steroid that represents the final product of the hypothalamic-pituitary-adrenal (HPA) axis stress response, to brain organoids in an in vitro model can result in changes in brain organoids through single-cell transcriptomics.

In this study, in addition to inflammatory markers such as C-reaction protein (CRP) and interleukin (IL)-6, which are routinely measured in clinical studies, we aimed to explore a broader range of biomarkers using WBC single-cell transcriptomics and plasma proteomics. Furthermore, we focused on identifying biomarkers in patients with MDD with atypical features accompanied by psychotic symptoms, an area that has been relatively under-researched.

## Methods and materials

### Participants

This study was approved by the Institutional Review Board of our University Hospital (2022-08-003). Korean women diagnosed with MDD with atypical features accompanied by psychotic symptoms such as IOR, were recruited as the patient group. Patients with psychotic symptoms may or may not meet criteria for psychotic features specifier of MDD. A trained psychiatrist interviewed the participants and made the psychiatric diagnoses. Healthy controls were recruited from among individuals without a psychiatric diagnosis, including those with mood disorders. Participants with neurological conditions, such as seizures, cerebrovascular disease, brain neoplasm, traumatic brain injury, or intellectual disabilities, were excluded from the study.

All individuals who agreed to participate underwent physical examination, which included measurements of height, weight, blood pressure, pulse, and body temperature, as well as the collection of psychiatric and medical histories. Self-reported measures were used to assess mood (HAM-D), anxiety (HAM-A), affect (PANAS), IOR (IOR subscale of the SPQ), perceived stress (visual scale), life events (LEC-5), problematic drinking (the third question of Alcohol Use Disorders Identification Test-Korean, AUDIT-K), and suicidal ideation (the ninth question of Patient Health Questionnaire-9, PHQ-9). We collected blood samples from the study participants, isolated WBCs, and performed WBC single-cell transcriptomics. Additionally, iPSC generation and WES were performed using samples from a single patient. Furthermore, blood tests were performed to obtain a complete blood count and differential count and to measure the levels of inflammatory markers such as CRP and IL-6.

### Statistical analyses

Statistical analyses were performed using R and Python (including scipy and pandas) software. For parametric variables that followed a normal distribution, the Student’s *t*-test was applied. The Mann-Whitney U test was used for non-normally distributed variables. The chi-square test was performed for nonparametric variables.

Further detailed methods are described in the **Supplementary Methods**.

## Results

### Higher stress levels, psychiatric symptoms, elevated WBC count, and increased CRP levels in the patient group

This study compared 7 women with atypical depression and psychotic symptoms to 10 healthy female controls (Table 1). The patient group was younger (29.1 years vs. 37.4 years), and only 1 of 10 patients was married, compared with 7 of 10 women in the control group. In terms of employment, 4 of 10 patients were employed, while all 10 participants in the control group were employed. Physical examination and social history revealed that the pulse rate and amount of smoking were significantly higher in the patient group.

**Table 1.** Clinical Characteristics and Laboratory Results of Participants.

|  | **Patients (n=7)** | **Controls (n=10)** | **Statistic** | **p** |
| --- | --- | --- | --- | --- |
| **Age, years** | 29.1 | 37.4 | -2.75 | 0.015 |
| **Married** | 1 (14.3%) | 7 (70.0%) | 3.14 | 0.076 |
| **Employment** | 2 (28.6%) | 9 (90.0%) | 4.38 | 0.036 |
| **SBP, mmHg** | 123.4 | 113.8 | 1.46 | 0.165 |
| **DBP, mmHg** | 79.1 | 71.8 | 1.73 | 0.105 |
| **PR, bpm** | 93.4 | 76.2 | 2.25 | 0.040 |
| **BW, kg** | 82.6 | 66.7 | 47 | 0.270 |
| **Height, cm** | 163.1 | 162.5 | 0.24 | 0.812 |
| **Smoking, amount** | 0.4 | 0.0 | 50 | 0.033 |
| **Alcohol, per week** | 0.1 | 0.4 | 29 | 0.469 |
| **Alcohol, amount** | 0.1 | 0.4 | 26 | 0.294 |
| **Problematic drinking** | 0.3 | 0.3 | 31 | 0.645 |
| **Self-rated Scales** |  |  |  |  |
| **Stress visual scale** | 62.0 | 33.3 | 2.71 | 0.016 |
| **Suicidal idea** | 1.0 | 0 | 60 | 0.003 |
| **spq_ior_total** | 4.6 | 1.6 | 68 | 0.001 |
| **LEC-5_num** | 4.3 | 1.2 | 3.01 | 0.009 |
| **PSS** | 26.7 | 16.0 | 3.81 | 0.002 |
| **HAM-D** | 22.7 | 1.1 | 70 | 0.001 |
| **HAM-A** | 25.4 | 1.4 | 70 | 0.001 |
| **Blood Tests** |  |  |  |  |
| **WBC, $\times 10^9/\text{mm}^3$** | 7.5 | 5.3 | 2.24 | 0.041 |
| **RBC, $\times 10^9/\text{mm}^3$** | 12.4 | 12.7 | 1.79 | 0.094 |
| **Hb, g/dL** | 12.4 | 12.7 | -0.32 | 0.757 |
| **Hct, %** | 38.8 | 38.9 | 40 | 0.660 |
| **MCV, fL** | 85.5 | 91.6 | 37 | 0.884 |
| **MCH, pg** | 27.5 | 30.0 | 30 | 0.669 |
| **MCHC, g/dL** | 31.9 | 61.8 | 25 | 0.378 |
| **RDW, fL** | 15.3 | 13.4 | 51 | 0.118 |
| **PLT, $\times 10^9/\text{mm}^3$** | 306.4 | 238.0 | 47 | 0.270 |
| **MPV, fL** | 8.6 | 9.0 | -0.66 | 0.518 |
| **Neutrophil, %** | 59.0 | 56.1 | 0.66 | 0.522 |
| **Lymphocyte, %** | 31.8 | 33.7 | -0.50 | 0.625 |
| **Monocyte, %** | 5.2 | 6.4 | -1.58 | 0.136 |
| **Eosinophil, %** | 2.4 | 2.6 | 35.5 | 1.000 |
| **Basophil, %** | 0.3 | 0.5 | -1.32 | 0.208 |
| **ANC, /$\mu\text{L}$** | 4519.4 | 3045.3 | 1.96 | 0.069 |
| **CRP ($> 0.3$), mg/L** | 3 (42.9%) | 0 (0%) | 4.63 | 0.031 |
| **HbA1c, %** | 12.3 | 5.2 | 35 | 1.000 |
| **IL-6** | 6.1 | 2.7 | 1.97 | 0.080 |

- - 1 For parametric variables, we performed Student's t-test or Mann-Whitney U test, and for non- 2 parametric variables, we performed the chi-square test. Abbreviations: ANC, absolute neutrophil count; BW, body weight; CRP, C-reactive protein; DBP, diastolic blood pressure; HAM- A, Hamilton anxiety rating scale; HAM-D, Hamilton depression rating scale; Hb, hemoglobin; HbA1c, hemoglobin A1c; Hct, hematocrit; IL-6; interleukin 6; MCH, mean corpuscular hemoglobin; MCHC, mean corpuscular hemoglobin concentration; MCV, mean corpuscular volume; MPV, mean platelet volume; PLT, platelet; PR, pulse rate; RBC, red blood cell; RDW, red cell distribution width; SBP, systolic blood pressure; WBC, white blood cell

The use of self-report questionnaires indicated that subjective stress perception, measured via the Stress Visual Scale, was higher in the patient group (62 points) than in the control group (33.3 points). Stress perception, as measured using the Perceived Stress Scale (PSS), was also elevated in the patient group. Suicidal ideation was more prevalent in the patient group. The severities of depressive symptoms (assessed using the Hamilton Depression Rating Scale, HAM-D) and anxiety symptoms (measured using the Hamilton Anxiety Rating Scale, HAM-A) were both higher in the patient group. Affective states, such as anger and nervousness (measured using the Positive and Negative Affect Scale, PANAS), also had higher scores in the patient group. Furthermore, women in the patient group had higher scores on the IOR subscale of the Schizotypal Personality Questionnaire (SPQ) than those in the control group. Traumatic events, assessed using LEC-5 (Life Events Checklist), were more frequent in the patient group (4.29 vs. 1.2 events).

Blood tests revealed that the patient group exhibited elevated WBC counts and a higher frequency of mild CRP elevation. There were no between-group differences in neutrophil and monocyte differential counts as well as IL-6 levels.

### Plasma protein expression changes in patients with atypical depression and psychotic symptoms

Plasma proteomic analysis was performed on 5 patients and 10 healthy controls using proximity extension assay (Fig. 1).

**Fig. 1.**
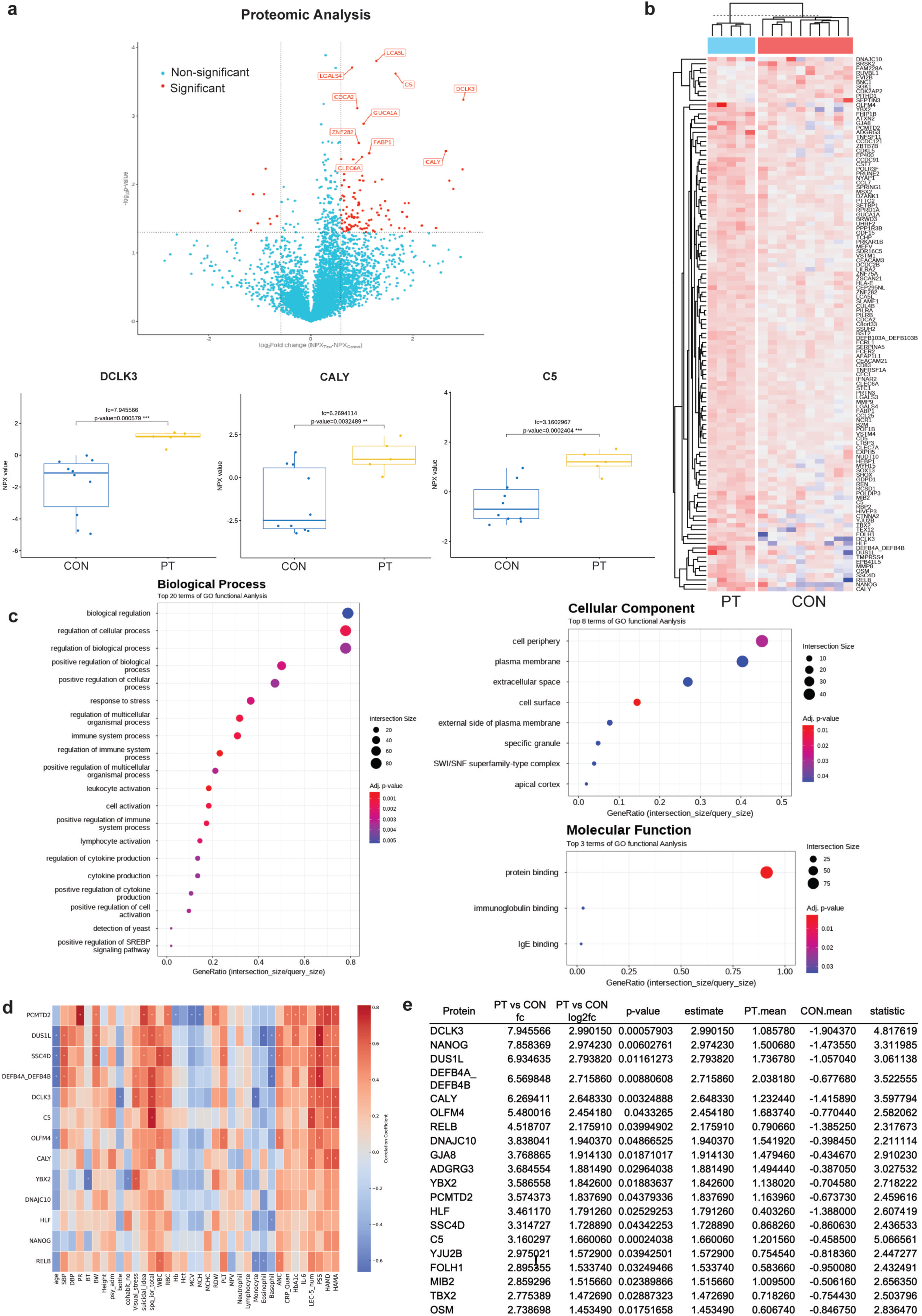
Plasma Proteomic Finding from Patients with Atypical Depression and Psychotic Symptoms. (a) A volcano plot showed increased expression of plasma proteins including DCLK3, CALY, C5 in the patient group (PT) compared to the control group (CON) (b) A heat map of the two-way hierarchical clustering to differentiate plasma protein expression between PT and CON (c) Gene ontology (GO) term analyses showed the most prominent changes in regulating cellular process in the biological process, cell periphery in the cellular component, and protein binding in the molecular function (d) Detailed list of differentially expressed proteins between PT and CON

In proteomic results, the highest level of expression increase was observed in DCLK3 (Doublecortin Like Kinase 3) (fold change, 7.95), a gene associated with synapse formation, followed by NANOG (Homeobox Transcription Factor NANOG) (7.86), which are embryonic and neural stem cell-related transcription factors, DUS1L (Dihydrouridine Synthase 1 Like) (6.93), and DEFB4A_DEFB4B (Defensin Beta 4A and 4B) (6.57). CALY (Calcyon Neuron-Specific Vesicular Protein) (6.27), also involved in nervous system functions such as synapse formation, showed similarly high expression. Immune-related genes including OLFM4 (Olfactomedin 4) (5.48), RELB (RELB Proto-Oncogene, NF-κB Subunit) (4.52), C5 (Complement Component 5) (3.16), and SSC4D (Scavenger Receptor Cysteine Rich Family Member with 4 Domains) (3.31) also exhibited notable increases. Additionally, genes such as DNAJC10 (DnaJ Heat Shock Protein Family Member C10) (3.84), YBX2 (Y-Box Binding Protein 2) (3.59), HLF (Hepatic Leukemia Factor) (3.46), and PCMTD2 (Protein-L-Isoaspartate (D-Aspartate) O-Methyltransferase Domain Containing 2) (3.57), which are involved in various cellular functions including transcription regulation, protein folding and degradation, and RNA binding, showed more than 3-fold increases in expression (Fig. 1a - b and e).

Gene ontology analysis revealed that significantly regulated genes were predominantly associated with molecular functions such as protein binding and immunoglobulin binding, cellular components including the plasma membrane and extracellular space, and biological processes related to immune responses, leukocyte activation, and cellular regulation (Fig. 1c).

Correlation analyses revealed that DCLK3 expression was positively associated with the Visual Stress Scale (r = 0.607), SPQ-IOR total score (r = 0.702), LEC-5 event count (r = 0.567), PSS (r = 0.608), HAM-D (r = 0.620), and HAM-A (r = 0.652). CALY expression correlated with the SPQ-IOR total score (r = 0.516), LEC-5 event count (r = 0.549), HAM-D (r = 0.620), and HAM-A (r = 0.644). C5 showed strong correlations with the SPQ-IOR total score (r = 0.779), LEC-5 event count (r = 0.699), PSS (r = 0.467), HAM-D (r = 0.591), and HAM-A (r = 0.646). Remarkably, DUS1L expression exhibited the strongest correlation overall, with PSS (r = 0.812) (Fig. 1e).

### WBC single-cell transcriptomic alteration in patients with atypical depression and psychotic symptoms

In this study, WBC single-cell RNA sequencing (scRNA-seq) was performed on 6 patients and 10 healthy controls, with data analyzed using the Seurat package^44^ (Fig. 2).

**Fig. 2.**
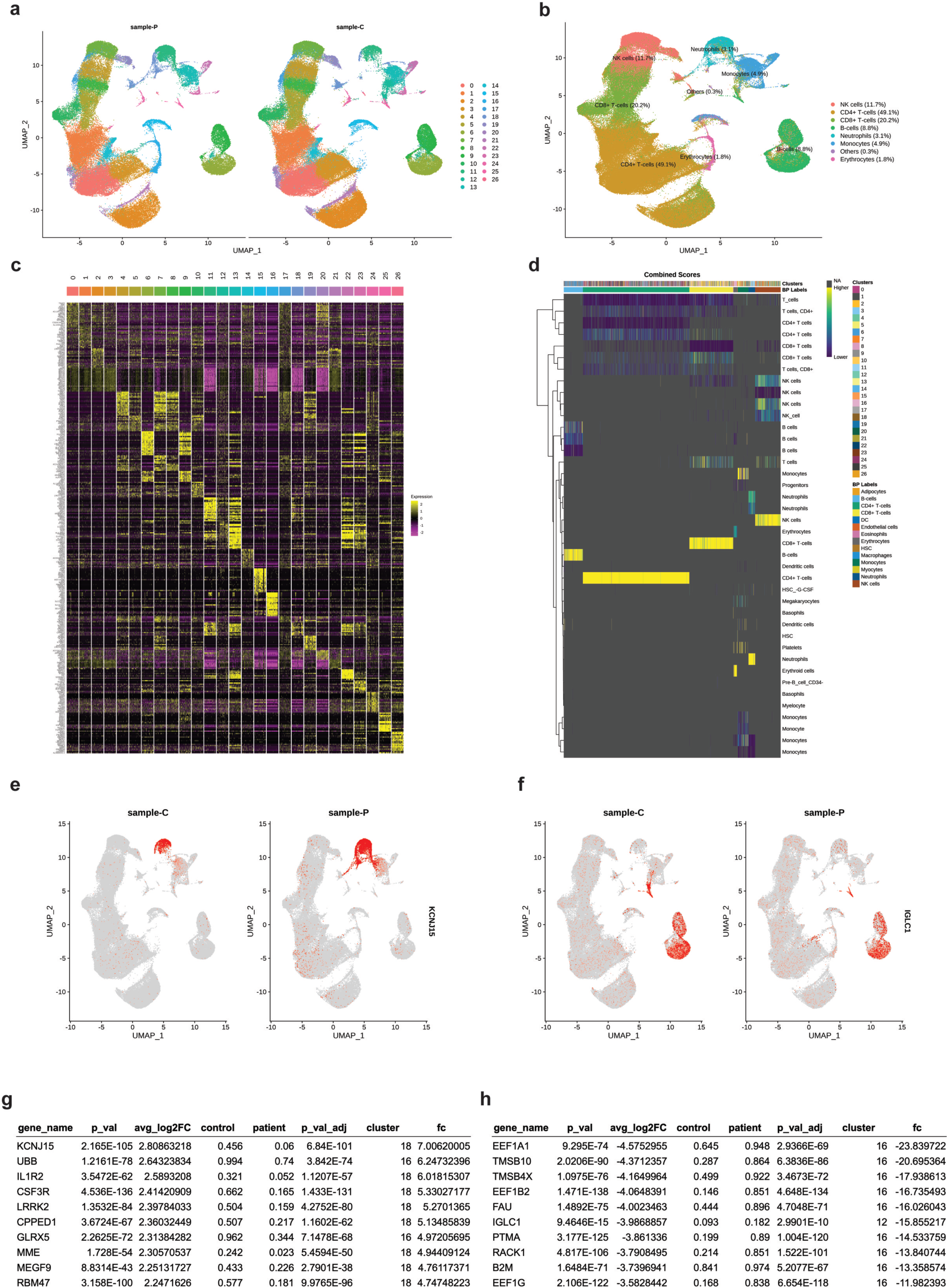
Differential WBC Single-cell Transcriptomic Profiles in the Patient Group Elevated Inflammatory Gene Expression in Neutrophils and Monocytes. (a-b) Identified cell types by Seurat analysis from single-cell RNA sequencing in WBC between the patient (sample-P) and healthy control (sample-C) groups. Graph theory-based cell type clustering of **B** is presented. Heatmap showing the gene expression patterns for the top genes in each cluster (c and d). (e and f) FeaturePlot showing differentially expressed genes between the patient and healthy control groups at the cell level for each cluster. The expression level of *KCNJ15* increased more than seven times (fc 7.01) in neutrophils of the patient group compared with those of the healthy control group (e). The expression level of *IGLC1* decreased more than 15 times (fc -15.86) in B cells of the patient group compared with those of the healthy control group (f). (g and h) List of the top 10 differentially expressed genes of single-cell RNA sequencing of WBCs between the patient and healthy control groups.

The WBC scRNA-seq results revealed notable transcriptomic differences between the groups. In cluster 18, classified as neutrophils, the patient group exhibited higher expressions of genes, such as *KCNJ15* (fold change, [fc] 7.01), *IL1R2* (fc 6.01), *CSF3R* (fc 5.33), *LRRK2* (fc 5.27), *CPPED1* (fc 5.13), *GLRX5* (fc 4.97), *MME* (fc 4.94), *MEGF9* (fc 4.76), and *RBM47* (fc 4.75). In cluster 16, classified as monocytes, *UBB* (fc 6.24) was also highly expressed in the patient group. Conversely, in cluster 16, decreased transcriptomic expression was observed in the patient group for genes, such as *EEF1A1* (fc 23.84), *TMSB10* (fc 20.70), *TMSB4X* (fc 17.94), *EEF1B2* (fc 16.74), *FAU* (fc 16.03), *PTMA* (fc 14.53), *RACK1* (fc 13.84), *B2M* (fc 13.36), and *EEF1G* (fc 11.98). In cluster 12, classified as B cells, *IGLC1* (fc 15.86) showed reduced expression in the patient group. Notably, WBC scRNA-seq findings did not align with the proteomic results.

### Growth and single-cell transcriptomic changes in brain organoids in a patient with atypical depression and psychotic symptoms

To investigate changes in patient brain organoids, monocytes from a patient were reprogrammed into iPSCs using the Sendai virus and subsequently differentiated into brain organoids. As a control, brain organoids were generated from iPSCs (obtained from adipose tissue-derived mesenchymal stem cells from a healthy female) provided by the National Stem Cell Bank in South Korea (Fig. 3 and Supplementary Fig. 1)^45^. These organoids were cultured for 60 days, followed by a 1-week treatment with DMX, a potent corticosteroid, to simulate stress-induced HPA axis activation (Fig. 3a). A size difference in the brain organoids between the patient and control groups became apparent, particularly on day 60 (Fig. 3b).

**Fig. 3.**
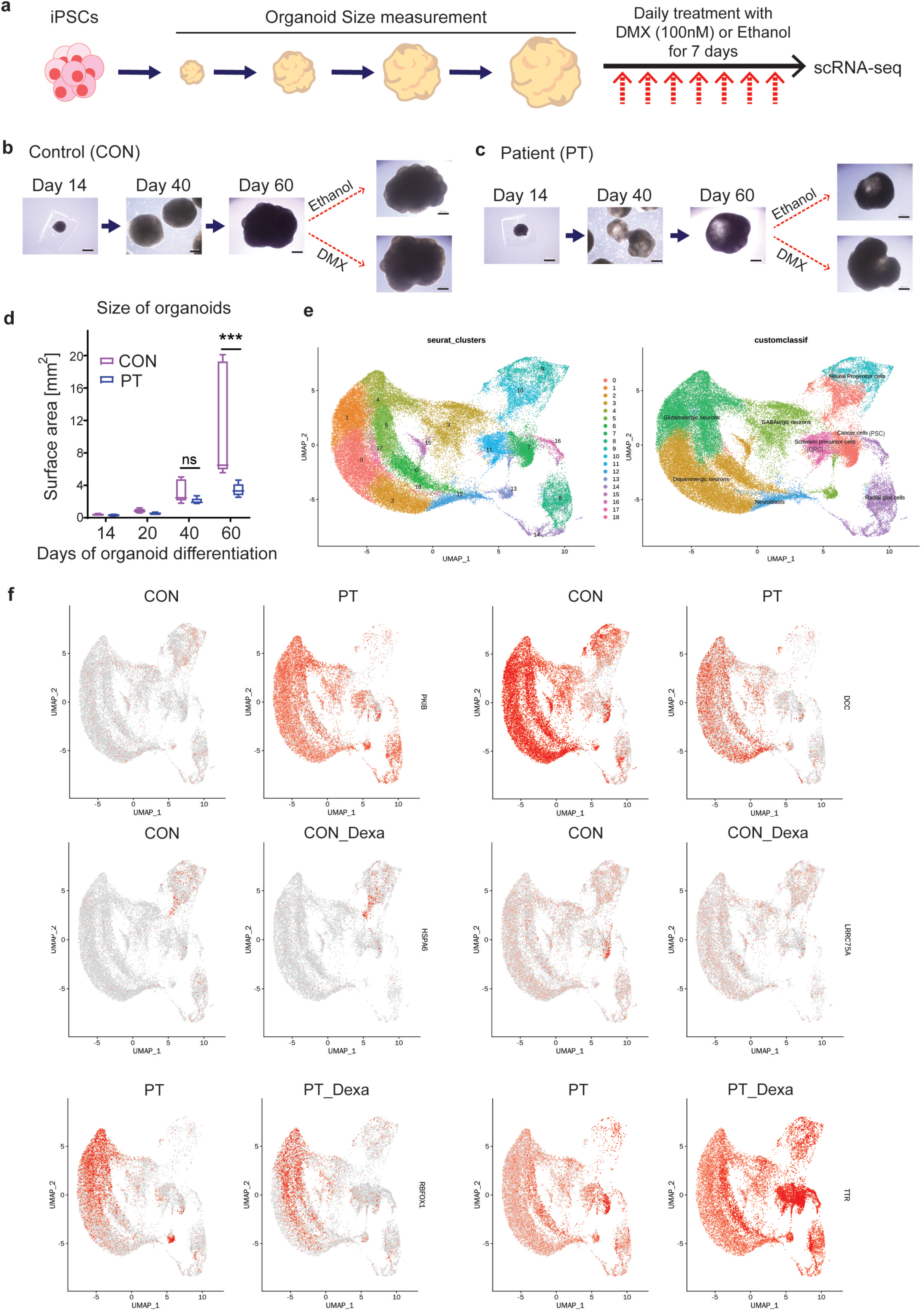
Transcriptomic Alterations in Brain Organoids Derived from a Patient with Atypical Depression and Psychotic Symptoms Increased Variability in Response to Dexamethasone Treatment. (a) Experimental scheme for single-cell RNA sequencing. Brain organoids were generated using induced pluripotent stem cells (iPSCs) derived from both a patient (PT) and a healthy control (CON). Organoids were cultured for 60 days. To simulate stress conditions, the organoids were treated with dexamethasone (DMX, 100 nM) or ethanol for 1 week. (b) Bright-field images showing CON brain organoids on days 14, 40, and 60, as well as after treatment with DMX or ethanol. Scale bar = 500 μm. (c) Bright-field images showing PT organoids on days 14, 40, and 60, as well as after treatment with DMX or ethanol. Scale bar = 500 μm. (d) The brain organoids derived from the patient group were smaller than those from the healthy control group on day 60. Student’s t-test (n = 5 CON, n = 5 PT) ***, *p*<0.001; ns, not significant. (e) Cell types identified by Seurat analysis of single-cell RNA sequencing of brain organoids. Graph theory-based cell type clustering is presented. (F) FeaturePlot showing differentially expressed genes between patients and healthy controls at the cellular level for each cluster. In the patient group, decreased expression of *DCC* was observed in clusters 13 (GABAergic neurons), 17 (glutamatergic neurons), 5 (glutamatergic neurons), 18 (neuroblasts), and 1 (glutamatergic neurons). In contrast, increased expression of *PKIB* was observed in clusters 4 (glutamatergic neurons), 15 (dopaminergic neurons), 5 (glutamatergic neurons), 1 (glutamatergic neurons), 8 (radial glial cells), 3 (GABAergic neurons), and 17 (glutamatergic neurons). When comparing the CON and CON_Dexa groups, *LRRC75A* expression was reduced, whereas *HSPA6* expression was increased in cluster 10 (cancer cells) in the CON_Dexa group. In the PT and PT_Dexa groups, *TTR* expression increased in clusters 10 (cancer cells), 13 (GABAergic neurons), 7 (cancer cells), 11 (Schwann precursor cells), 8 (radial glial cells), 16 (radial glial cells), 14 (radial glial cells), and 15 (dopaminergic cells), whereas decreased expression was observed for *RBFOX1* in cluster 13 (GABAergic cells).

scRNA-seq results for the five organoids from each group revealed significant transcriptomic differences (Supplementary Fig. 2 and Supplementary Table 1). In the patient group, decreased expression was observed in genes such as *DCC* (clusters 13 [GABAergic neurons], 17 [glutamatergic neurons], 5 [glutamatergic neurons], 18 [neuroblast], and 1 [glutamatergic neurons]), *PMCH* (cluster 9 [neural progenitor cells]), *HTR2C* (cluster 7 [cancer cell]), and *TTR* (cluster 7 [cancer cell]), and *DPP10* (cluster 4 [glutamatergic neurons]). The following genes were highly expressed: *PKIB* (clusters 4 [glutamatergic neurons], 15 [dopaminergic neurons], 5 [glutamatergic neurons], 1 [glutamatergic neurons], 8 [radial glial cells], 3 [GABAergic neurons], and 17 [glutamatergic neurons]), *KCNMB2* (cluster 13 [GABAergic neurons]), *TSHZ2* (cluster 13 [GABAergic neurons]), and *CSMD1* (cluster 4 [glutamatergic neurons]).

When comparing brain organoids of controls before (CON) and after (CON_Dexa) DMX treatment, *LRRC75A* expression was reduced by fc 2.70 in the CON_Dexa group, while *HSPA6* (cluster 10 [cancer cells]) and *PMCH* (cluster 9 [Schwann precursor cells]) expressions increased by fc 2.24 and fc 2.23, respectively.

An analysis of brain organoids of patients without (PT) and with (PT_Dexa) DMX treatment (Supplementary Table 2) demonstrated increased expressions of *TTR* (clusters 10 [cancer cells], 13 [GABAergic neurons], 7 [cancer cells], 11 [Schwann precursor cells], 8 [radial glial cells], 16 [radial glial cells], 14 [radial glial cells], and 15 [dopaminergic cells]), *PMCH* (cluster 9 [neural precursor cells]), and *CHL1* (cluster 15 [dopaminergic cells]), with decreased expressions of *RBFOX1* (cluster 13 [GABAergic cells]), *NMB* (cluster 9 [neural precursor cells]), *LRRC75A* (cluster 7 [cancer cells]), *PTPRD, ANKS1B, KCNH7, KAZN, CCSER1, DLGAP1*, and *IGFBP5* (cluster 10 [cancer cells]).

Compared with the CON group, which showed two increased and one decreased transcriptomic changes (Supplementary Table 3), the PT group exhibited a greater response to DMX treatment, with 23 increased and 34 decreased transcriptomic changes, suggesting a more significant transcriptomic variability in the PT_Dexa group than in the CON_Dexa group.

In comparison with proteomic findings, *CALY* expression was significantly reduced in the PT group (PT) relative to the CON group with fc −1.12 and −1.13 (clusters 1 [glutamatergic neurons] and 4 [glutamatergic neurons]). *DCLK3* expression was also decreased in the PT group (fc −1.09, cluster 2 [dopaminergic neurons]). Notably, *DCLK3* expression was upregulated following DMX treatment (PT_Dexa vs. PT), with an FC of 1.09 in the same cluster.

### Genomic changes in a patient with atypical depression and psychotic symptoms

To further investigate genetic polymorphisms in a patient who generated the brain organoid and not included in the proteomic analysis, we performed whole-exome sequencing (WES) at 300x coverage using WBC of the patient (Supplemenatry Fig. 3).

In total, 90,174 single-nucleotide polymorphisms (SNPs) and insertions/deletions were identified. The results were analyzed by focusing on 12,868 genes with a quality control score > 30 and those classified as having a moderate or higher predicted impact; we observed a heterozygous SNP in the *COMT* gene (chromosome 22, rs4680, c.472 G>A), which is known as a schizophrenia risk gene. Another heterozygous SNP was identified in rs6267 (c.214 G>T), which is related to drug response. In addition, a missense variant in *CALY* (c.589 G>A, Ala197Thr) was identified in parallel proteomic analysis.

Other notable findings included *PTEN* (rs34003473, c.1321-3dupT), a gene associated with neurogenesis; *PARK2* (rs1801474, c.500 G>A), linked to Parkinson’s disease; and *GIGYF2* (rs371622656, c.3692_3693insGC). Immune-related genes such as *HLA-A*, *HLA-B*, *HLA-DQB1*, *HLA-DRB1*, and *HLA-DRB5* were also identified; however, these were classified as benign variants (Supplementary Table 5).

Interestingly, although *COMT* expression showed no significant differences between the groups in WBC scRNA-seq, brain organoid scRNA-seq revealed slightly reduced expression in the patient group, specifically in clusters 0 (dopaminergic neurons, fc 1.07), 1 (glutamatergic neurons, fc 1.08), 8 (radial glial cells, fc 1.17), and 9 (neural precursor cells, fc 1.60).

## Discussion

In this study, we combined clinical data, WBC scRNA-seq, plasma proteomics, brain organoids, and WES to investigate the transcriptomic and genomic changes in patients with MDD with atypical features accompanied by psychotic symptoms. Our findings provide new insights into the immunological and neurological changes in this patient population, as well as the potential molecular mechanisms underlying the disease course. This is the first study to implement a precision medicine approach by integrating WBC single-cell transcriptomics and brain organoids with clinical data from patients with MDD.

The patient group exhibited higher levels of stress, trauma exposure, anxiety, and depression, with more frequent experiences of anger and nervousness. A previous study reported that 98% of patients with SMI had a history of lifetime traumatic events^46^. Although approximately 50% of the general population also reports lifetime traumatic events, the prevalence is significantly higher in individuals with SMI; traumatic events are well-known contributors to the onset and exacerbation of psychiatric disorders^47^. Given that the patient group had atypical depression with psychotic symptoms, we expected a higher prevalence of severe symptoms, social impairment, and pronounced biological abnormalities than other depressive disorders.

Although this was a cross-sectional study, we observed a higher prevalence of traumatic events, severe psychiatric symptoms, and increased immune responses in the patient group. Individuals who have experienced traumatic events, including those with post-traumatic stress disorder, reportedly exhibit increased inflammation and changes in immune cell profiles^47,48^. Childhood adversity also affects the immune system in adulthood^49^. Increased WBC count suggests an increased inflammatory response, indicating evidence of previous immune dysfunction in patients with mood disorders^50–53^. These results included stress-related immune dysfunction and neurological alterations, as evidenced by multi-omics approach including plasma proteomics, WBC scRNA-seq and brain organoid analyses in our study.

Among the proteins that showed significant alterations in plasma proteomics, DCLK3, CALY, and C5 have previously been implicated in psychiatric and neurological disorders. DCLK3 is a neuron-specific kinase predominantly expressed in the striatum and hippocampus, and is suggested to contribute to the neurobiological underpinnings of psychiatric disorders through its involvement in the regulation of anxiety-related behaviors and spatial memory ^54,55^. Moreover, DCLK3 has been implicated in brain transcriptomic association studies of schizophrenia and bipolar affective disorder ^56^.

CALY, a protein involved in presynaptic vesicle formation, has been reported in patients with attention-deficit/hyperactivity disorder (ADHD), schizophrenia, and cocaine use disorder ^57,58^. C5, a complement protein, has been found to be elevated in the cerebrospinal fluid (CSF) of patients with schizophrenia and major depressive disorder ^59^, and its expression has been negatively correlated with the thickness of the superior frontal gyrus in individuals with schizophrenia ^60^.

The WBC scRNA-seq results were particularly notable in neutrophils and monocytes, with the latter having a potential central nervous system involvement. Genes such as *IL1R2*^61^ and *Csf3R*^62^, which are key molecules in the inflammatory response, were highly expressed in neutrophils, suggesting elevated inflammation in the patient group. Additionally, *LRRK2* associated with Parkinson’s disease, was upregulated and is also known to regulate microglial inflammation^63^. *Ubb*^64^ was highly expressed in monocytes and played a crucial role in antigen presentation and immune regulation. The downregulation of *B2m*^65^ in neutrophils, which is involved in MHC class I, suggests alterations in immune signaling. Previous studies on MDD have shown limited findings in WBC scRNA-seq. One study reported increased inflammatory and immune-metabolic pathways in monocytes of patients with high CRP levels^66^. Although our study did not detect an increase in MMP8 levels, as seen in stress-vulnerable mice^67^, our results align with those of existing studies on immune dysregulation in MDD.

In brain organoid experiments, patient blood-derived organoids showed reduced growth by day 60 compared with those of healthy controls. Transcriptomic analysis revealed changes from immature neural stem cells to mature neurons. Key genes, such as *DCC* (neural migration and axon guidance^68^), *PMCH* (melanin-concentrating hormone^69^), and *HTR2C* (serotonin 2C receptor associated with stress response^70^), were differentially expressed, highlighting disruptions in neural development and signaling. Following DMX treatment, genes involved in the cellular stress response, such as *LRRC75A*^71^ and *HSPA6* ^72^, showed altered expression in the controls, whereas the patient group exhibited increased expressions of *TTR*^73^*, PMCH, and CHL1*^74^, which are all associated with brain development. In comparison with the findings of Cruceanu et al.^75^, our study utilized DMX to replicate HPA-axis activation and observed greater transcriptomic changes in the patient group, indicating heightened brain vulnerability.

Given the scarcity of brain organoid research in MDD, our study offers a pioneering approach that combines WBC scRNA-seq and brain organoid models to explore personalized medicine for psychiatric disorders. Direct comparative studies on the prevalence of somatic mutations in brain organoids derived from patient groups have not yet been conducted ^39^. Notably, recent research on brain somatic mutations in schizophrenia patients has identified brain-specific somatic genetic variants in postmortem brain tissues, primarily affecting genes that play critical roles in neural signaling and development ^76^. In our study, we performed WES on brain organoids to compare conditions before and after stress simulation using DMX treatment (Supplementary Fig – WES organoid?). These findings provide direct evidence of brain somatic mutations induced by stress and highlight their potential role in severe mental illnesses, which is often associated with significant life adversity and traumatic events. This underscores the importance of further research into the role of brain somatic mutations in the pathogenesis of severe mental illnesses.

Our WES analysis from blood of the patient identified well-known genetic polymorphisms related to brain organoid changes and symptoms. Notably, the identification of a heterozygous SNP in the *COMT* gene (rs4680), linked to schizophrenia risk^77^, provided insights into the psychotic symptoms observed in the patient group. Polymorphisms in genes associated with neurogenesis (*PTEN*^78^) and neurodegenerative disorders (*PARK2*^79^) were also observed, explaining the impaired brain organoid growth. Interestingly, although *COMT* expression differences were not significant in WBC scRNA-seq, they were evident in brain organoids, highlighting the distinct expression profiles of immune and neural tissues. In our study, the *COMT* rs4680 G>A polymorphism was identified; moreover, and the *COMT* rs4680 A allele is reportedly associated with a better therapeutic response to antipsychotics, such as olanzapine^80–82^, and increased suicidal attempts^83^. This suggests that incorporating WES data during patient treatment could help predict the treatment response, potentially leading to earlier symptom remission and faster functional recovery. Additionally, we propose that future studies should investigate the effects of pharmacological treatments on brain organoids under DMX exposure, with the goal of establishing a platform for evaluating drug efficacy in the context of HPA axis activation as a model of stress.

In scRNA-seq analysis of patient-derived brain organoids, *CALY* expression was significantly reduced compared to controls. Similarly, *DCLK3* expression was downregulated in the same samples. Notably, *DCLK3* expression was restored following DMX treatment in patient-derived organoids. Among peripheral measurements, plasma proteomics more closely mirrored the molecular alterations observed in brain organoids than did WBC scRNA-seq. These findings suggest that plasma proteomics may capture peripheral signatures of subtle brain pathophysiology, including changes associated with psychiatric conditions such as MDD. Although WBC scRNA-seq was less reflective of neural alterations, it may still offer complementary insight into immune system activity that could modulate brain function. Given its higher concordance with brain organoid transcriptomic changes, plasma proteomics may serve as a primary approach for psychiatric biomarker discovery, with WBC scRNA-seq contributing additional context to refine molecular profiling.

The limitations of our study include the small cohort focused on patients with MDD with psychotic symptoms, which may not be generalizable across all patients with MDD. Additionally, validation techniques, such as quantitative polymerase chain reaction, were not performed for the WBC and brain organoid scRNA-seq data.

Moreover, brain organoids cannot include microglia because they are composed solely of cells of an ectodermal origin. This limitation highlights the importance of using WBC scRNA-seq to indirectly assess the state of microglia, for example through monocytes, which served as a complementary approach in our study. Future studies should focus on incorporating models that reflect or include microglia. However, the strength of our study lies in its comprehensive approach of integrating clinical data, plasma protein expression, immune profiling, and brain–organoid research to investigate the pathophysiology of MDD. This pioneering work sheds light on immune and neurological abnormalities in MDD and provides valuable insights for future research on precision medicine in psychiatry.

In conclusion, this study provides a pioneering approach by multi-omics, integrating WBC scRNA-seq, plasma proteomics, brain organoid scRNA-seq, WES, and clinical data to explore the biological underpinnings of MDD with atypical features accompanied by psychotic symptoms. Our findings revealed immune and neuronal dysregulation - particularly in neutrophils, monocytes, and plasma proteins - and identified significant transcriptomic changes in brain organoids under stressful conditions. These results suggest a heightened vulnerability to stress in patients with MDD and emphasize the importance of immune-neuronal interactions in the pathophysiology of the disease. Our work provides valuable insights for advancing precision medicine in psychiatry.

## Author contributions

JH and YK contributed to the conception and design of the study; SC and SHC contributed to the acquisition of clinical information and data; IA performed iPSC and brain organoid experiments; JH and YK wrote the manuscript and prepared the figures.

## Funding

This work was supported by the National Research Foundation of Korea (NRF) grant funded by the Korean government (MSIT) (NRF-2021R1C1C1003266 to Y.K.; RS-2024-00335144 and RS-2024-00398786 to J.H.), and the Korea Health Technology R&D Project through the Korea Health Industry Development Institute (KHIDI) funded by the Ministry of Health & Welfare, Republic of Korea (HI22C0492 and RS-2024-00438635 to Y.K.). I.A. was supported by the NRF through the Basic Science Research Program (RS-2024-00410041).

## Conflict of interest

There is no conflict of interest.

## Notes

### Competing Interest Statement

The authors have declared no competing interest.

