## supplementary tables and figures, methods for "Exploration of Novel Biomarkers through a Precision Medicine Approach Using Multi-omics and Brain Organoids in Patients with Atypical Depression and Psychotic Symptoms"

**Method and Materials**

**Single cell RNA sequencing (scRNA-seq) – WBC and brain organoids**

Briefly, WBCs were isolated from whole blood of study participants, and dead cells were removed for WBC scRNA-seq. Five brain organoids per sample, differentiated for 67 days, were dissociated, and dead cells were removed for brain organoid scRNA-seq. cDNA libraries targeting 10,000 cells per sample were prepared using the 10x Chromium Single Cell 3’ Protocol (10x Genomics, CA, USA). The prepared libraries were sequenced on an Illumina HiSeq X system (Illumina, CA, USA).

For bioinformatic analysis, raw base call files were converted to FASTQ format using ‘mkfastq,’ and sorting, filtering, barcode counting, and UMI counting were performed with ‘cellranger.’ Subsequent analyses, including data preprocessing, quality control, dimensionality reduction, graph-based cell clustering, identification of top-expressed biomarkers, gene set enrichment (using gProfiler), differential gene expression between groups, cell subtype classification, and population analysis, were conducted using the Seurat 4.0.5 R package. Cell type classification was based on the method described by Zeisel et al. Single-cell RNA sequencing and bioinformatics analyses were performed by Macrogen (Seoul, South Korea). Mitochondrial, ribosomal, RBC-related and MALAT1 genes/isoforms were excluded from the analysis.

**Whole Exome Sequencing**

The Twist NGS Target Enrichment workflow utilizes ultra-long 120-mer biotinylated cDNA baits to capture and enrich regions of interest from genomic fragment libraries. gDNA libraries were prepared using 50 ng of input DNA and the Twist Library Preparation EF Kit, with TruSeq-compatible Y-adapters (Illumina, CA, USA). DNA quantity and quality were assessed using PicoGreen and agarose gel electrophoresis. Libraries were fragmented to 200 bp, followed by end-repair, ‘A’ tailing, adapter ligation, and PCR amplification. Final products were quantified using TapeStation DNA screentape D1000 (Agilent, CA, USA) and PicoGreen. For exome capture, 1500 ng of indexed libraries were pooled and hybridized with Twist exome probes, followed by washing, amplification, and final quantification using qPCR. Sequencing was performed on an Illumina platform, utilizing the Sequencing-by-Synthesis method for high accuracy.

For bioinformatic analysis, raw images and base calling were generated using Illumina’s RTA software and converted into FASTQ format with bcl2fastq v2.20.0, with demultiplexing set to zero barcode mismatches. Sequencing quality was assessed using FastQC. Paired-end reads were mapped to the human reference genome using BWA-MEM, producing BAM files without unordered sequences or alternate haplotypes. PCR duplicates were removed using MarkDuplicates from the Picard-tools package. BAM files were further processed with Base Quality Score Recalibration (BQSR) to adjust for sequencing errors. Variant genotyping was performed with HaplotypeCaller (GATK), and SNPs/indels were identified. Variants were filtered using GATK’s VariantFiltration tool and annotated with SnpEff, incorporating data from dbSNP, 1000 Genomes, and other databases. The final output was in VCF format with additional annotations from ESP6500, ClinVar, dbNSFP, and ACMG.

**Reprogramming hiPSCs**

Four days prior to the introduction of reprogramming factors (referred to as day -4)**,** peripheral blood mononuclear cells (PBMCs) were collected according to the following procedure. A patient’s blood sample, collected in BD Vacutainer K2 EDTA (K2E) blood collection tubes, was diluted in 10 ml of DPBS (Welgene, LB201-02) and gently layered over 20 ml of lymphocyte separation medium (Corning 25-072-CV). The sample was centrifuged at 300×g for 20 minutes at room temperature (RT). After centrifugation, the middle layer containing PBMCs was carefully transferred to a 50 ml tube (SPL 50450). The cells were then washed twice with DPBS, followed by centrifugation at 300×g for 8 minutes each time at RT. Three million PBMCs were seeded into a 24-well plate (SPL 30024) at a density of 0.5 million cells per well and cultured in Stempro-34 medium (Gibco 10639-011) supplemented with 100 ng/ml SCF (Gibco PHC2111), 100 ng/ml FLT-3 (Gibco PHC9414), 20 ng/ml IL-3 (Gibco PHC0034), and 20 ng/ml IL-6 (Gibco PHC0065), referred to as the complete PBMC medium. The cells were cultured with half media changes every other day for 4 days.

On day 0, 0.5 million PBMCs were placed in a round-bottom tube and treated with Sendai viral particles using the CytoTune-iPS Sendai Reprogramming Kit (Gibco A16518), according to the manufacturer's instructions, at multiplicities of infection (MOIs) of KOS = 5, hc-Myc = 5, and hKlf4 = 3. The round-bottom tube containing the PBMC and virus mixture was centrifuged at 1000×g for 30 minutes at RT. After centrifugation, the PBMCs were resuspended in complete PBMC medium and transferred to a single well of a 12-well plate (SPL 30012). On day 1, cells were collected into a 15 ml tube (SPL 50215) and centrifuged at 200×g for 10 minutes at RT to remove Sendai viruses, then resuspended in complete PBMC medium in a 24-well plate and cultured for 2 days without media changes. On day 3, the reprogrammed cells were plated on mouse embryonic fibroblasts (MEFs), which had been inactivated with 10 μg/ml of mitomycin C (AG Scientific M-1108) for 2 hours. The Institutional Review Board of KAIST has approved the protocols for using MEFs in accordance with the relevant ethical standards and regulations (KH2021-069). From day 4 to day 6, half of the medium was changed with StemPro-34. On day 7, half of the medium was replaced with iPSC medium consisting of DMEM/F12 (Gibco 12400024) supplemented with 1% MEM non-essential amino acids (NEAA; Gibco 11140050), 14.3 mM sodium bicarbonate (Sigma S5761), 1 mM L-glutamine (Sigma S5792), 0.1 mM β-Mercaptoethanol (2-ME, Merck M3148), 20% knockout serum replacement (KSR, Gibco 10828028), and 10 ng/ml bFGF (R&D Systems 4114-TC). On day 8, a full media change with the same iPSC medium was performed. The reprogrammed iPSC colonies were monitored under a microscope until they emerged. The generated iPSCs were maintained on MEFs in iPSC medium and passaged onto new plates every 6 days using collagenase (Gibco 17104019).

**iPSC validation:** **Live cell imaging, Reverse transcription (RT)-PCR, Karyotyping**

TRA1-60 was visualized using the TRA-1-60 Alexa Fluor™ 488 Conjugate Kit for Live Cell Imaging (ThermoFisher A25618) according to the manufacturer’s instructions. Briefly, the molecular probe was centrifuged at 10,000×g for 2 minutes and then applied to the prepared iPSCs at a 1:50 ratio in culture media. The cells were incubated at 37°C for 30 minutes. After incubation, the cells were washed 2-3 times with FluoroBrite™ DMEM (ThermoFisher A1896701) and imaged using the Olympus IX71 microscope.

To validate the expression of genes associated with pluripotent stem cells, *KLF4*, *SOX2*, *OCT4* and *NANOG* were amplified by RT-PCR. Total RNA was first extracted under RNase-free conditions using TRIzol (Invitrogen 15596-018), chloroform (Junsei 28560-0350), isopropanol (Merck 109634.1011), and ethanol (Merck 100983.1011). cDNA was synthesized from the total RNA using oligo dT and the RevertAid First Strand cDNA synthesis kit (ThermoFisher K1621) according to the manufacturer’s instructions. PCR was then performed using the cDNA, Ex Taq (Takara R001A) and the following primers: KLF4_Fwd 5’ CGCTGGCGGGAGGAGCTCTC 3’, KLF4_Rev 5’ GGTGACAGTCCCTGCTGCTC 3’, SOX2_Fwd 5’ CCGTTCATCGACGAGGCTAA 3’, SOX2_Rev 5’ ATTGGTGTTCTCTTTTGCAG 3’, OCT4_Fwd 5’ CTTGGAGACCTCTCAGCCTG3’, OCT4_Rev 5’ AGCAGGGCTGGATGCCTTCA 3’, NANOG_Fwd 5’ GATACTCATAAAGCCGCTAC 3’, NANOG _Rev 5’ CCCCACTTGCTCATTCCCAA 3’, ACTB_Fwd 5’ CGAGCACGGCATCGTCACCAA 3’, ACTB _Rev 5’GCAGCACGGGGTGCTCCTCG 3’. PCR was performed using a Bio-RAD C-1000 Touch Thermal Cycler under the following conditions: denaturation at 95°C for 1 minute, annealing at 55°C to 65°C for 1 minute, and elongation at 72°C for 1 to 2 minutes, for a total of 30 amplification cycles.

Karyotyping analysis was performed by Dx&Vx, Inc. (Seoul, South Korea).

**Forebrain organoids differentiation**

On day 0, iPSCs were dissociated using collagenase and transferred to non-adherent dishes (SPL 10060) with media (1), containing 2 nM Dorsomorphin (STEMCELL Technologies 72102), 2 nM A-83 (STEMCELL Technologies 72022), 1% MEM NEAA, 0.1 mM 2-ME, 20% KSR, and 1% penicillin/streptomycin (PS) (Welgene LS202-02) in DMEM F-12/Glutamax (Gibco 10565042). A full media change was performed on day 1, and half media changes were done on days 3 and 4. On days 5 and 6, the organoids were transitioned to media (2), which contained 1 nM CHIR-99021 (STEMCELL Technologies 72052), 0.1 nM SB-431242 (Cayman CAY-13031), 1% MEM NEAA, 1% PS, and N2 supplement (Gibco 17502048) in DMEM F-12/Glutamax, with half media changes. On day 7, 6 organoids were embedded in 30 µl of Matrigel (Corning 354230) and incubated at 37°C, 5% CO2 for 40 minutes. The Matrigel-coated organoids were transferred to media (2) and cultured with half media changes every other day until day 13. On day 14, embedded organoids were dissected out of the Matrigel using a surgical scalpel (Jeungdo bio H-2201-3). The separated organoids were cultured on a shaker (N-BIOTEK 101SRC) rotating continously at 80 rpm in media (3), containing N2 and B27 supplement (Gibco12587010), 0.1 mM 2-ME, 1% PS, and 2.5 ng/ml insulin (Sigma I9278). Thereafter, the organoids were cultured with half media changes every other day. For the DMX (Sigma D4902) treatment, 100 nM DMX diluted in ethanol was added to the culture media daily for one week, starting on day 61 after the organoids were differentiated for 60 days. As a control, 100% ethanol was added to the organoids daily for one week, starting on day 61.

**Supplementary Tables**

**Supplementary Table 1.** List of the top 20 differentially expressed genes of single-cell RNA sequencing of brain organoids between a patient and a healthy control, sorted by fold change (fc)

Down

| **gene_name** | **p_val** | **avg_log2FC** | **PT** | **CON** | **p_val_adj** | **cluster** | **fc** |
| --- | --- | --- | --- | --- | --- | --- | --- |
| DCC | 1.4369E-54 | -3.28 | 0.133 | 0.82 | 4.6132E-50 | 13 | -9.73 |
| PMCH | 2.2159E-16 | -2.96 | 0.173 | 0.475 | 7.1142E-12 | 9 | -7.79 |
| DCC | 8.0442E-33 | -2.63 | 0.228 | 0.918 | 2.5827E-28 | 17 | -6.20 |
| HTR2C | 1.0157E-21 | -2.50 | 0.004 | 0.164 | 3.2609E-17 | 7 | -5.66 |
| DCC | 2.604E-245 | -2.46 | 0.324 | 0.963 | 8.359E-241 | 5 | -5.51 |
| DCC | 5.2132E-31 | -2.45 | 0.635 | 1 | 1.6738E-26 | 18 | -5.48 |
| DCC | <2.225e-308 | -2.36 | 0.407 | 0.993 | <2.225e-308 | 1 | -5.14 |
| TTR | 2.6654E-40 | -2.36 | 0.581 | 0.824 | 8.5574E-36 | 7 | -5.13 |
| DCC | 7.413E-308 | -2.34 | 0.362 | 0.986 | 2.38E-303 | 4 | -5.07 |
| DPP10 | 1.465E-230 | -2.32 | 0.183 | 0.816 | 4.705E-226 | 4 | -4.99 |
| HTR2C | 6.5454E-19 | -2.30 | 0.087 | 0.582 | 2.1015E-14 | 10 | -4.92 |
| DCC | <2.225e-308 | -2.27 | 0.369 | 0.984 | <2.225e-308 | 0 | -4.82 |
| RSPO3 | 0.00027995 | -2.21 | 0.011 | 0.16 | 1 | 16 | -4.63 |
| UNC5D | 6.8663E-68 | -2.19 | 0.007 | 0.324 | 2.2045E-63 | 6 | -4.57 |
| DCC | 1.068E-168 | -2.19 | 0.335 | 0.936 | 3.427E-164 | 6 | -4.56 |
| DPP10 | 2.3542E-26 | -2.17 | 0.071 | 0.467 | 7.5585E-22 | 13 | -4.49 |
| TCF7L2 | 1.9103E-23 | -2.14 | 0.077 | 0.434 | 6.1334E-19 | 13 | -4.40 |
| UNC5D | 1.283E-252 | -2.07 | 0.014 | 0.546 | 4.119E-248 | 2 | -4.21 |
| DCC | <2.225e-308 | -2.06 | 0.373 | 0.99 | <2.225e-308 | 2 | -4.16 |
| AL157778.1 | 1.5951E-17 | -2.03 | 0.172 | 0.707 | 5.1212E-13 | 15 | -4.08 |

Up

| **gene_name** | **p_val** | **avg_log2FC** | **PT** | **CON** | **p_val_adj** | **cluster** | **fc** |
| --- | --- | --- | --- | --- | --- | --- | --- |
| AC069277.1 | 4.9481E-88 | 2.84 | 0.849 | 0.003 | 1.5886E-83 | 14 | 7.17 |
| AC069277.1 | 1.396E-259 | 2.84 | 0.795 | 0.01 | 4.483E-255 | 8 | 7.15 |
| PKIB | 5.671E-278 | 2.34 | 0.796 | 0.067 | 1.821E-273 | 4 | 5.08 |
| AC069277.1 | 1.266E-128 | 2.28 | 0.489 | 0.004 | 4.063E-124 | 6 | 4.86 |
| AC069277.1 | 5.511E-135 | 2.23 | 0.684 | 0.005 | 1.769E-130 | 12 | 4.69 |
| PKIB | 3.4662E-43 | 2.13 | 0.736 | 0.053 | 1.1129E-38 | 15 | 4.37 |
| PKIB | 6.421E-172 | 2.05 | 0.582 | 0.012 | 2.061E-167 | 5 | 4.15 |
| PKIB | <2.225e-308 | 2.04 | 0.809 | 0.033 | <2.225e-308 | 1 | 4.12 |
| KCNMB2 | 1.1361E-07 | 2.03 | 0.515 | 0.303 | 0.0036477 | 13 | 4.09 |
| HES5 | 0.00564366 | 2.01 | 0.269 | 0.111 | 1 | 16 | 4.04 |
| PKIB | 5.921E-183 | 2.01 | 0.701 | 0.07 | 1.901E-178 | 8 | 4.02 |
| PKIB | 6.017E-254 | 1.92 | 0.614 | 0.065 | 1.932E-249 | 3 | 3.79 |
| PKIB | 3.054E-21 | 1.92 | 0.691 | 0 | 9.8052E-17 | 17 | 3.79 |
| AC069277.1 | <2.225e-308 | 1.91 | 0.679 | 0.005 | <2.225e-308 | 2 | 3.77 |
| TSHZ2 | 1.2636E-15 | 1.91 | 0.638 | 0.279 | 4.057E-11 | 13 | 3.76 |
| CSMD1 | 1.0364E-76 | 1.89 | 0.521 | 0.19 | 3.3274E-72 | 4 | 3.70 |
| PKIB | 1.0726E-30 | 1.86 | 0.889 | 0.085 | 3.4437E-26 | 18 | 3.63 |
| PKIB | <2.225e-308 | 1.83 | 0.763 | 0.036 | <2.225e-308 | 0 | 3.57 |
| AC069277.1 | 1.7765E-18 | 1.83 | 0.459 | 0.016 | 5.7037E-14 | 13 | 3.55 |
| PKIB | 3.693E-91 | 1.81 | 0.417 | 0.027 | 1.1857E-86 | 6 | 3.49 |

**Supplementary Table 2.** List of the top 20 differentially expressed genes of single-cell RNA sequencing of brain organoids between a healthy control with or without dexamethasone treatment, sorted by fold change (fc)

Down

| **gene_name** | **p_val** | **avg_log2FC** | **PT_Dexa** | **PT** | **p_val_adj** | **cluster** | **fc** |
| --- | --- | --- | --- | --- | --- | --- | --- |
| AL589740.1 | 0.00018711 | -2.03 | 0.031 | 0.369 | 1 | 13 | -4.09 |
| PBX3 | 0.0001835 | -1.93 | 0.188 | 0.549 | 1 | 13 | -3.80 |
| ERBB4 | 7.8895E-05 | -1.93 | 0.375 | 0.689 | 1 | 13 | -3.80 |
| TENM3 | 0.01557005 | -1.91 | 0.312 | 0.508 | 1 | 13 | -3.75 |
| TCF7L2 | 0.00030282 | -1.87 | 0.094 | 0.434 | 1 | 13 | -3.65 |
| TAC1 | 0.00304307 | -1.77 | 0.125 | 0.393 | 1 | 13 | -3.41 |
| CRABP1 | 0.00889225 | -1.73 | 0.125 | 0.344 | 1 | 13 | -3.31 |
| AC096570.1 | 0.01809003 | -1.50 | 0.094 | 0.295 | 1 | 13 | -2.82 |
| LRRC75A | 9.4454E-17 | -1.43 | 0.037 | 0.163 | 3.0326E-12 | 7 | -2.70 |
| CNTN5 | 0.05928509 | -1.41 | 0.125 | 0.279 | 1 | 13 | -2.67 |
| HIST1H4C | 0.04458496 | -1.32 | 0.306 | 0.481 | 1 | 16 | -2.50 |
| NLGN1 | 0.00077345 | -1.27 | 0.061 | 0.321 | 1 | 16 | -2.41 |
| TTR | 0.03097761 | -1.24 | 0.719 | 0.918 | 1 | 13 | -2.36 |
| SSR2 | 8.329E-05 | -1.23 | 0.204 | 0.58 | 1 | 16 | -2.35 |
| EDIL3 | 0.00021438 | -1.22 | 0.125 | 0.484 | 1 | 13 | -2.33 |
| KLHL13 | 0.0019165 | -1.20 | 0.094 | 0.369 | 1 | 13 | -2.30 |
| SLC4A7 | 0.03323071 | -1.16 | 0.082 | 0.222 | 1 | 16 | -2.23 |
| WLS | 0.0003104 | -1.14 | 0.125 | 0.484 | 1 | 13 | -2.21 |
| UBE2C | 0.01298131 | -1.14 | 0.041 | 0.198 | 1 | 16 | -2.21 |
| HECW1 | 0.00167105 | -1.14 | 0.02 | 0.222 | 1 | 16 | -2.20 |

Up

| **gene_name** | **p_val** | **avg_log2FC** | **PT_Dexa** | **PT** | **p_val_adj** | **cluster** | **fc** |
| --- | --- | --- | --- | --- | --- | --- | --- |
| TTR | 0.0177317 | 2.59 | 0.8 | 0.932 | 1 | 14 | 6.03 |
| SPARCL1 | 0.22980245 | 2.00 | 0.102 | 0.049 | 1 | 16 | 4.00 |
| IGFBP5 | 0.09041268 | 1.59 | 0.245 | 0.148 | 1 | 16 | 3.02 |
| PRR16 | 0.00740844 | 1.55 | 0.188 | 0.049 | 1 | 13 | 2.93 |
| TEX14 | 0.25759235 | 1.38 | 0.062 | 0.148 | 1 | 13 | 2.60 |
| LMO4 | 0.89028444 | 1.31 | 0.25 | 0.287 | 1 | 13 | 2.48 |
| BCAN | 0.0334408 | 1.29 | 0.224 | 0.099 | 1 | 16 | 2.44 |
| AC012409.2 | 8.5944E-05 | 1.28 | 0.531 | 0.213 | 1 | 13 | 2.43 |
| SORCS3 | 0.03124673 | 1.28 | 0.375 | 0.221 | 1 | 13 | 2.42 |
| SYT11 | 0.08056664 | 1.27 | 0.306 | 0.21 | 1 | 16 | 2.40 |
| LINGO2 | 0.00029866 | 1.20 | 0.625 | 0.352 | 1 | 13 | 2.30 |
| HTR2C | 0.81049557 | 1.20 | 0.189 | 0.188 | 1 | 14 | 2.29 |
| GPC5 | 0.01946949 | 1.18 | 0.344 | 0.18 | 1 | 13 | 2.26 |
| ZFYVE16 | 0.00147781 | 1.17 | 0.286 | 0.086 | 1 | 16 | 2.25 |
| HSPA6 | 4.5903E-07 | 1.16 | 0.19 | 0.122 | 0.01473748 | 10 | 2.24 |
| PMCH | 3.0689E-26 | 1.16 | 0.648 | 0.475 | 9.853E-22 | 9 | 2.23 |
| LHFPL3 | 0.96996457 | 1.16 | 0.375 | 0.41 | 1 | 13 | 2.23 |
| RAB21 | 0.10368809 | 1.14 | 0.122 | 0.049 | 1 | 16 | 2.21 |
| ATCAY | 0.19920939 | 1.13 | 0.163 | 0.099 | 1 | 16 | 2.20 |
| SLIT2 | 0.0621193 | 1.13 | 0.406 | 0.279 | 1 | 13 | 2.19 |

**Supplementary Table 3.** List of the top 20 differentially expressed genes of single-cell RNA sequencing of brain organoids between a patient with or without dexamethasone treatment, sorted by fold change (fc)

Down

| **gene_name** | **p_val** | **avg_log2FC** | **CON_Dexa** | **CON** | **p_val_adj** | **cluster** | **fc** |
| --- | --- | --- | --- | --- | --- | --- | --- |
| RBFOX1 | 2.0127E-37 | -2.55 | 0.101 | 0.48 | 6.462E-33 | 13 | -5.85 |
| NFIA-AS2 | 1.6178E-37 | -2.00 | 0.043 | 0.221 | 5.1943E-33 | 7 | -3.99 |
| XIST | 2.2604E-35 | -1.71 | 0.007 | 0.175 | 7.2571E-31 | 6 | -3.28 |
| HES5 | 0.03833708 | -1.67 | 0.192 | 0.269 | 1 | 16 | -3.19 |
| XIST | 3.2075E-32 | -1.54 | 0.012 | 0.274 | 1.0298E-27 | 12 | -2.90 |
| AL157778.1 | 9.9533E-36 | -1.51 | 0.037 | 0.385 | 3.1956E-31 | 13 | -2.84 |
| XIST | 2.4125E-18 | -1.45 | 0.041 | 0.36 | 7.7457E-14 | 14 | -2.74 |
| NMB | 5.4988E-12 | -1.40 | 0.37 | 0.578 | 1.7655E-07 | 9 | -2.65 |
| XIST | 5.9374E-71 | -1.40 | 0.018 | 0.301 | 1.9063E-66 | 8 | -2.63 |
| XIST | 1.6072E-36 | -1.37 | 0.007 | 0.156 | 5.1602E-32 | 5 | -2.58 |
| LRRC75A | 8.8719E-20 | -1.34 | 0.032 | 0.137 | 2.8484E-15 | 7 | -2.54 |
| PTPRD | 2.4752E-26 | -1.31 | 0.141 | 0.337 | 7.947E-22 | 7 | -2.48 |
| ZFPM2-AS1 | 2.7089E-25 | -1.28 | 0.054 | 0.202 | 8.6973E-21 | 7 | -2.43 |
| LINC01456 | 2.3221E-39 | -1.27 | 0.108 | 0.523 | 7.4552E-35 | 13 | -2.41 |
| IL1RAPL1 | 5.9289E-06 | -1.24 | 0.128 | 0.291 | 0.19035313 | 10 | -2.37 |
| ANKS1B | 2.3116E-13 | -1.24 | 0.088 | 0.199 | 7.4215E-09 | 7 | -2.36 |
| KCNH7 | 5.6155E-17 | -1.24 | 0.109 | 0.249 | 1.8029E-12 | 7 | -2.35 |
| KAZN | 1.6176E-13 | -1.23 | 0.135 | 0.262 | 5.1934E-09 | 7 | -2.35 |
| FTX | 2.0207E-19 | -1.20 | 0.158 | 0.324 | 6.4878E-15 | 7 | -2.30 |
| CCSER1 | 8.7584E-21 | -1.19 | 0.135 | 0.305 | 2.812E-16 | 7 | -2.29 |

Up

| **gene_name** | **p_val** | **avg_log2FC** | **pct.1** | **pct.2** | **p_val_adj** | **cluster** | **fc** |
| --- | --- | --- | --- | --- | --- | --- | --- |
| TTR | 1.5336E-34 | 2.62 | 0.992 | 0.612 | 4.9238E-30 | 10 | 6.15 |
| TTR | 1.3393E-90 | 2.44 | 0.993 | 0.597 | 4.3001E-86 | 13 | 5.44 |
| PMCH | 5.4692E-08 | 2.37 | 0.372 | 0.173 | 0.00175595 | 9 | 5.17 |
| TTR | 4.747E-164 | 2.24 | 0.98 | 0.581 | 1.524E-159 | 7 | 4.71 |
| TTR | 3.197E-194 | 2.19 | 0.987 | 0.602 | 1.026E-189 | 11 | 4.57 |
| TTR | 3.263E-199 | 2.03 | 0.985 | 0.601 | 1.047E-194 | 8 | 4.10 |
| TTR | 1.1605E-30 | 2.03 | 0.993 | 0.516 | 3.726E-26 | 16 | 4.09 |
| TTR | 6.5753E-56 | 1.95 | 0.986 | 0.647 | 2.1111E-51 | 14 | 3.85 |
| CHL1 | 8.2962E-23 | 1.91 | 0.817 | 0.08 | 2.6636E-18 | 15 | 3.75 |
| TTR | 2.809E-24 | 1.82 | 1 | 0.621 | 9.0187E-20 | 15 | 3.53 |
| TTR | 2.686E-244 | 1.77 | 0.992 | 0.6 | 8.625E-240 | 3 | 3.40 |
| TTR | 2.142E-114 | 1.73 | 0.988 | 0.605 | 6.878E-110 | 12 | 3.31 |
| TTR | 2.159E-190 | 1.72 | 0.988 | 0.607 | 6.931E-186 | 5 | 3.30 |
| TTR | 4.502E-156 | 1.69 | 0.983 | 0.595 | 1.445E-151 | 6 | 3.23 |
| TTR | 2.6777E-40 | 1.67 | 1 | 0.589 | 8.5969E-36 | 17 | 3.18 |
| CCN2 | 5.8358E-07 | 1.62 | 0.291 | 0.058 | 0.01873649 | 10 | 3.08 |
| TTR | <2.225e-308 | 1.61 | 0.992 | 0.595 | <2.225e-308 | 2 | 3.06 |
| GULP1 | 1.6423E-13 | 1.60 | 0.581 | 0.194 | 5.2729E-09 | 10 | 3.04 |
| TTR | 6.58E-264 | 1.53 | 0.993 | 0.621 | 2.113E-259 | 4 | 2.89 |
| HTR2C | 1.6697E-09 | 1.47 | 0.41 | 0.087 | 5.3606E-05 | 10 | 2.78 |

**Supplementary Table 4.** High putative impact results from whole exome sequencing of patient-derived and control-derived brain organoids

Control

| **CHROM** | **POS** | **REF** | **ALT** | **DP** | **AD** | **QUAL** | **MQ** | **Zygosity** | **FILTER** | **Effect** | **Gene_Name** |
| --- | --- | --- | --- | --- | --- | --- | --- | --- | --- | --- | --- |
| chr1 | 8324503 | C | CG | 65 | 10 | 162.73 | 60 | HET | PASS | frameshift_variant | SLC45A1 |
| chr1 | 33013330 | G | C | 116 | 10 | 12.05 | 57 | HET | MG_SNP_Filter | structural_interaction_variant | AK2 |
| chr1 | 2.13E+08 | G | T | 75 | 17 | 68.77 | 60 | HET | MG_SNP_Filter | splice_acceptor_variant&intron_variant | RPS6KC1 |
| chr1 | 2.4E+08 | C | CA | 76 | 5 | 40.73 | 58.43 | HET | MG_INDEL_Filter | frameshift_variant | FMN2 |
| chr1 | 2.4E+08 | CA | C | 72 | 5 | 43.73 | 58.35 | HET | MG_INDEL_Filter | frameshift_variant | FMN2 |
| chr2 | 9849441 | T | TA | 115 | 19 | 196.73 | 60 | HET | PASS | frameshift_variant | TAF1B |
| chr2 | 32141890 | A | AT | 113 | 14 | 20.77 | 60 | HET | MG_INDEL_Filter | splice_acceptor_variant&intron_variant | SPAST |
| chr2 | 97185336 | GC | G | 89 | 8 | 45.73 | 43.36 | HET | MG_INDEL_Filter | frameshift_variant | ANKRD36 |
| chr2 | 97185340 | C | G | 89 | 8 | 54.77 | 43.28 | HET | MG_SNP_Filter | stop_gained | ANKRD36 |
| chr2 | 97185341 | A | AG | 88 | 8 | 45.73 | 43.17 | HET | MG_INDEL_Filter | frameshift_variant | ANKRD36 |
| chr2 | 1.3E+08 | G | A | 141 | 15 | 55.77 | 58.61 | HET | MG_SNP_Filter | stop_gained | SMPD4 |
| chr2 | 1.97E+08 | C | CA | 108 | 18 | 167.73 | 60 | HET | PASS | frameshift_variant | CCDC150 |
| chr3 | 32704796 | G | GT | 52 | 11 | 106.73 | 60 | HET | PASS | splice_acceptor_variant&intron_variant | CNOT10 |
| chr3 | 1.14E+08 | G | GT | 105 | 13 | 41.73 | 60 | HET | MG_INDEL_Filter | frameshift_variant | USF3 |
| chr4 | 53453080 | CAG | C | 88 | 12 | 147.73 | 60 | HET | MG_INDEL_Filter | frameshift_variant | FIP1L1 |
| chr5 | 42808306 | CG | C | 116 | 13 | 195.73 | 60 | HET | MG_INDEL_Filter | frameshift_variant | SEPP1 |
| chr5 | 42808309 | T | TC | 115 | 13 | 177.73 | 60 | HET | MG_INDEL_Filter | frameshift_variant | SEPP1 |
| chr5 | 42808316 | AG | A | 113 | 9 | 63.73 | 60 | HET | MG_INDEL_Filter | frameshift_variant | SEPP1 |
| chr5 | 1.76E+08 | G | GT | 98 | 10 | 12.96 | 60 | HET | MG_INDEL_Filter | frameshift_variant&stop_lost | ARL10 |
| chr6 | 32584366 | C | T | 53 | 9 | 249.77 | 52.13 | HET | PASS | stop_gained | HLA-DRB1 |
| chr6 | 80007991 | A | ACCCC | 117 | 16 | 57.73 | 60 | HET | MG_INDEL_Filter | frameshift_variant | TTK |
| chr6 | 1.59E+08 | G | C | 97 | 17 | 18.82 | 60 | HET | MG_SNP_Filter | splice_donor_variant&intron_variant | FNDC1 |
| chr7 | 1.01E+08 | CCGGT | C | 73 | 7 | 54.73 | 59.11 | HET | MG_INDEL_Filter | frameshift_variant | MUC3A |
| chr7 | 1.01E+08 | T | TGAA | 71 | 6 | 24.78 | 59.08 | HET | MG_INDEL_Filter | stop_gained&disruptive_inframe_insertion | MUC3A |
| chr7 | 1.01E+08 | C | CT | 70 | 5 | 21.76 | 59.07 | HET | MG_INDEL_Filter | frameshift_variant | MUC3A |
| chr7 | 1.44E+08 | A | ATGGAGGCTGAGGAGGCCCAGCG | 151 | 9 | 562.73 | 25.66 | HET | PASS | frameshift_variant&stop_gained | ARHGEF5 |
| chr8 | 1.01E+08 | T | TC | 177 | 16 | 227.73 | 56.13 | HET | MG_INDEL_Filter | frameshift_variant | PABPC1 |
| chr8 | 1.01E+08 | T | C | 194 | 16 | 176.77 | 56.48 | HET | MG_SNP_Filter | structural_interaction_variant | PABPC1 |
| chr9 | 1.31E+08 | C | CT | 28 | 7 | 27.74 | 60 | HET | MG_INDEL_Filter | splice_acceptor_variant&intron_variant | NUP214 |
| chr10 | 12198692 | G | GT | 119 | 17 | 114.73 | 60 | HET | MG_INDEL_Filter | splice_acceptor_variant&intron_variant | CDC123 |
| chr10 | 1.25E+08 | G | A | 161 | 25 | 565.77 | 55.6 | HET | PASS | structural_interaction_variant | CTBP2 |
| chr10 | 1.25E+08 | G | A | 161 | 26 | 613.77 | 55.14 | HET | PASS | structural_interaction_variant | CTBP2 |
| chr10 | 1.25E+08 | C | T | 168 | 22 | 264.77 | 54.28 | HET | PASS | structural_interaction_variant | CTBP2 |
| chr10 | 1.25E+08 | G | A | 222 | 54 | 1829.77 | 53.05 | HET | PASS | structural_interaction_variant | CTBP2 |
| chr10 | 1.25E+08 | GCAAGTAGGGGTCATAAAATATGACGCTGAATC | G | 254 | 52 | 1704.73 | 53.78 | HET | PASS | structural_interaction_variant | CTBP2 |
| chr10 | 1.25E+08 | C | CCGAAGCCGATGAGGCCCAATATCTCCCCACGAATGCGGGCCTTTACTGAGGCCACCTCGCCAAT | 126 | 38 | 1371.73 | 55.52 | HET | PASS | splice_acceptor_variant&intron_variant | CTBP2 |
| chr11 | 1185057 | TGG | T | 211 | 25 | 488.73 | 57.45 | HET | PASS | frameshift_variant | MUC5AC |
| chr11 | 1185062 | A | AAC | 208 | 25 | 492.73 | 57.38 | HET | PASS | frameshift_variant | MUC5AC |
| chr11 | 63382198 | C | CA | 98 | 18 | 172.77 | 60 | HET | PASS | frameshift_variant | SLC22A9 |
| chr11 | 1.08E+08 | C | CT | 56 | 15 | 209.73 | 60 | HET | PASS | splice_acceptor_variant&intron_variant | ATM |
| chr12 | 11267457 | TGGACGAGGTGGGGGACCTTGGGACTGGTTTCCTCCTTGTGGGGGTGGTCCTTCTGGCTTTCCTGGACGAGGTGGGGGACCTTGAGGTTTGTTGCCTCCTTGTGGGGGTGGTCCTTCTGGCTTTCCC | T | 103 | 78 | 2595.73 | 51.85 | HET | PASS | frameshift_variant&splice_acceptor_variant&splice_region_variant&intron_variant | PRB3 |
| chr12 | 1.04E+08 | T | TATTG | 78 | 7 | 34.73 | 57.42 | HET | MG_INDEL_Filter | splice_donor_variant&intron_variant | TDG |
| chr12 | 1.04E+08 | T | C | 73 | 3 | 44.77 | 58.56 | HET | MG_SNP_Filter | splice_donor_variant&intron_variant | TDG |
| chr12 | 1.04E+08 | C | CT | 64 | 9 | 25.74 | 60 | HET | MG_INDEL_Filter | splice_acceptor_variant&intron_variant | HCFC2 |
| chr14 | 67864961 | C | CTTTTTT | 21 | 20 | 674.74 | 60 | HOM | PASS | splice_acceptor_variant&intron_variant | RAD51B |
| chr14 | 73105898 | AAG | A | 142 | 11 | 41.77 | 60 | HET | MG_INDEL_Filter | frameshift_variant | RBM25 |
| chr15 | 83008518 | CA | C | 43 | 10 | 94.73 | 60 | HET | PASS | frameshift_variant | C15orf40 |
| chr16 | 22008408 | G | GAGCAC | 162 | 45 | 1513.73 | 58.71 | HET | PASS | splice_donor_variant&intron_variant | C16orf52 |
| chr16 | 22075699 | A | ATGATCCTTTTCTGTATGGCTGCCCTAATATTTCCAATAG | 140 | 53 | 1933.73 | 55.87 | HET | PASS | splice_donor_variant&intron_variant | C16orf52 |
| chr16 | 81032554 | G | GT | 71 | 14 | 145.73 | 60 | HET | PASS | splice_acceptor_variant&intron_variant | CENPN |
| chr16 | 85071765 | C | CTT | 44 | 10 | 137.73 | 60 | HET | PASS | splice_acceptor_variant&intron_variant | KIAA0513 |
| chr19 | 4682866 | CAG | C | 213 | 20 | 191.73 | 60 | HET | MG_INDEL_Filter | frameshift_variant | DPP9-AS1 |
| chr19 | 8888859 | ACT | A | 148 | 13 | 95.73 | 57.4 | HET | MG_INDEL_Filter | frameshift_variant | MUC16 |
| chr19 | 17286682 | GTGTGTGTGTT | G | 18 | 7 | 250.73 | 60 | HET | PASS | frameshift_variant | ANKLE1 |
| chr19 | 32999679 | G | A | 141 | 14 | 13.95 | 46.75 | HET | MG_SNP_Filter | stop_gained | RHPN2 |
| chr19 | 52384820 | G | GGATCATGAGGTCAGGAGATC | 133 | 12 | 134.73 | 59.84 | HET | MG_INDEL_Filter | frameshift_variant&stop_gained | ZNF880 |
| chr19 | 56193031 | A | AG | 93 | 9 | 102.73 | 58.95 | HET | MG_INDEL_Filter | frameshift_variant | ZSCAN5B |
| chr19 | 56193035 | TG | T | 88 | 9 | 99.73 | 59.11 | HET | MG_INDEL_Filter | frameshift_variant | ZSCAN5B |
| chr20 | 46174847 | CG | C | 119 | 15 | 121.73 | 60 | HET | MG_INDEL_Filter | frameshift_variant | CDH22 |
| chr20 | 46174849 | CCCGAG | C | 126 | 15 | 100.73 | 60 | HET | MG_INDEL_Filter | frameshift_variant | CDH22 |
| chr21 | 28966883 | AT | A | 119 | 14 | 44.73 | 60 | HET | MG_INDEL_Filter | frameshift_variant | LTN1 |

Patient

| **CHROM** | **POS** | **REF** | **ALT** | **DP** | **AD** | **QUAL** | **MQ** | **Zygosity** | **FILTER** | **Effect** | **Gene_Name** |
| --- | --- | --- | --- | --- | --- | --- | --- | --- | --- | --- | --- |
| chr1 | 1.52E+08 | AT | A | 95 | 71 | 1828.73 | 60 | HOM | PASS | frameshift_variant&start_lost | HRNR |
| chr1 | 2.07E+08 | T | TG | 74 | 16 | 208.73 | 60 | HET | PASS | frameshift_variant | MAPKAPK2 |
| chr1 | 2.48E+08 | C | CTG | 73 | 7 | 45.73 | 52.21 | HET | MG_INDEL_Filter | frameshift_variant&stop_gained | OR2T2 |
| chr1 | 2.48E+08 | C | T | 90 | 13 | 81.77 | 31.71 | HET | MG_SNP_Filter | stop_gained | OR2T5 |
| chr1 | 2.49E+08 | A | ACG | 3 | 3 | 98.25 | 30.13 | HOM | PASS | frameshift_variant | OR2T29 |
| chr3 | 49721936 | T | TGCCCCCCC | 97 | 16 | 188.73 | 60 | HET | PASS | frameshift_variant | GMPPB |
| chr3 | 49721944 | CAG | C | 94 | 5 | 11.08 | 60 | HET | MG_INDEL_Filter | frameshift_variant | GMPPB |
| chr3 | 75738661 | C | CT | 129 | 12 | 97.73 | 55.67 | HET | MG_INDEL_Filter | frameshift_variant | ZNF717 |
| chr3 | 1.31E+08 | CT | C | 129 | 21 | 189.73 | 60 | HET | MG_INDEL_Filter | frameshift_variant | ASTE1 |
| chr3 | 1.96E+08 | C | CT | 124 | 16 | 84.73 | 60 | HET | MG_INDEL_Filter | frameshift_variant | PPP1R2 |
| chr4 | 38014723 | G | GCCGC | 96 | 17 | 245.73 | 60 | HET | PASS | frameshift_variant | TBC1D1 |
| chr4 | 38014733 | CAA | C | 89 | 11 | 103.73 | 60 | HET | MG_INDEL_Filter | frameshift_variant | TBC1D1 |
| chr4 | 1.4E+08 | TGCTGCTGCTGCTGC | T | 162 | 13 | 5605.73 | 59.98 | HET | PASS | frameshift_variant | MAML3 |
| chr4 | 1.4E+08 | TGCTGCTGCTGCTGC | TTGC | 162 | 112 | 5605.73 | 59.98 | HET | PASS | frameshift_variant | MAML3 |
| chr4 | 1.59E+08 | G | GCCCCCCC | 55 | 14 | 307.73 | 60 | HET | PASS | frameshift_variant | RAPGEF2 |
| chr5 | 1.16E+08 | T | TATATATATATATATGGAACTAAGACTATTACTTTGGAAAAGACCTG | 213 | 18 | 191.73 | 60 | HET | MG_INDEL_Filter | frameshift_variant&stop_gained | LVRN |
| chr5 | 1.19E+08 | C | CT | 61 | 11 | 72.73 | 60 | HET | MG_INDEL_Filter | splice_acceptor_variant&intron_variant | TNFAIP8 |
| chr5 | 1.73E+08 | A | AT | 78 | 9 | 122.73 | 60 | HET | PASS | splice_acceptor_variant&intron_variant | BNIP1 |
| chr6 | 32584178 | G | GCT | 76 | 25 | 888.73 | 40.08 | HET | PASS | frameshift_variant | HLA-DRB1 |
| chr6 | 32584183 | TGC | T | 75 | 24 | 918.73 | 40.08 | HET | PASS | frameshift_variant | HLA-DRB1 |
| chr6 | 32589642 | C | G | 105 | 16 | 434.77 | 46.22 | HET | PASS | splice_donor_variant&intron_variant | HLA-DRB1 |
| chr7 | 741431 | T | TGC | 65 | 15 | 246.73 | 60 | HET | PASS | frameshift_variant | DNAAF5 |
| chr7 | 1.42E+08 | A | G | 159 | 14 | 28.77 | 57.49 | HET | MG_SNP_Filter | structural_interaction_variant | MGAM |
| chr8 | 99832366 | TAG | T | 43 | 11 | 289.73 | 60 | HET | PASS | splice_acceptor_variant&intron_variant | VPS13B |
| chr8 | 99832368 | G | GTTTTTTTTTTTTTTTTTTT | 43 | 11 | 290.73 | 60 | HET | PASS | frameshift_variant&splice_region_variant | VPS13B |
| chr8 | 1.07E+08 | TA | T | 88 | 11 | 33.73 | 60 | HET | MG_INDEL_Filter | splice_acceptor_variant&intron_variant | OXR1 |
| chr8 | 1.44E+08 | T | TTCCC | 69 | 13 | 184.73 | 60 | HET | PASS | frameshift_variant | HSF1 |
| chr9 | 68303273 | A | T | 3 | 2 | 23.33 | 30.03 | HET | MG_SNP_Filter | stop_gained | FOXD4L3 |
| chr9 | 1.05E+08 | TGTTA | T | 148 | 14 | 150.73 | 51.35 | HET | MG_INDEL_Filter | frameshift_variant | OR13C5 |
| chr9 | 1.2E+08 | CT | C | 125 | 20 | 209.73 | 60 | HET | PASS | splice_acceptor_variant&intron_variant | CDK5RAP2 |
| chr10 | 86654816 | CAG | C | 82 | 10 | 30.73 | 60 | HET | MG_INDEL_Filter | frameshift_variant | OPN4 |
| chr10 | 1.25E+08 | T | G | 82 | 22 | 592.77 | 53.97 | HET | PASS | structural_interaction_variant | CTBP2 |
| chr11 | 1017036 | G | GCA | 471 | 34 | 116.73 | 54.3 | HET | MG_INDEL_Filter | frameshift_variant | MUC6 |
| chr11 | 64265028 | CAG | C | 75 | 9 | 52.73 | 60 | HET | MG_INDEL_Filter | frameshift_variant | PLCB3 |
| chr11 | 64265033 | CACTGGATGCCT | C | 78 | 7 | 45.73 | 60 | HET | MG_INDEL_Filter | frameshift_variant | PLCB3 |
| chr11 | 1.17E+08 | G | GA | 119 | 14 | 19.78 | 60 | HET | MG_INDEL_Filter | frameshift_variant | CEP164 |
| chr12 | 8932622 | C | T | 134 | 23 | 298.77 | 40.9 | HET | PASS | stop_gained | PHC1 |
| chr12 | 11091468 | A | ATT | 175 | 17 | 278.73 | 54.04 | HET | MG_INDEL_Filter | frameshift_variant | TAS2R43 |
| chr12 | 11091471 | TCC | T | 183 | 20 | 290.73 | 54.48 | HET | MG_INDEL_Filter | frameshift_variant | TAS2R43 |
| chr12 | 11308599 | GCCTCCTTGTGGGGGTGGTCTTTCTGGCTTTCCTGGAGGAGGTGGGGTACCTTGGGACTGGTTT | G | 208 | 27 | 500.73 | 58.7 | HET | PASS | frameshift_variant&splice_donor_variant&splice_region_variant&intron_variant | PRB4 |
| chr12 | 52949347 | G | GA | 89 | 9 | 90.73 | 57.97 | HET | MG_INDEL_Filter | frameshift_variant | KRT18 |
| chr12 | 1.04E+08 | A | ATT | 165 | 34 | 988.73 | 58.32 | HET | PASS | splice_donor_variant&intron_variant | TDG |
| chr12 | 1.04E+08 | G | GAGAGCGTGGAGT | 156 | 30 | 989.73 | 58.34 | HET | PASS | splice_donor_variant&intron_variant | TDG |
| chr12 | 1.22E+08 | CACCA | C | 47 | 8 | 77.73 | 60 | HET | MG_INDEL_Filter | frameshift_variant | SETD1B |
| chr13 | 21155151 | C | CAGTTTTCTT | 156 | 34 | 979.73 | 60 | HET | PASS | splice_acceptor_variant&intron_variant | SKA3 |
| chr13 | 21159987 | T | TCA | 115 | 29 | 927.73 | 60 | HET | PASS | splice_acceptor_variant&intron_variant | SKA3 |
| chr13 | 42159244 | C | CTTTTTTTTTTTTTTTTTTTTT | 25 | 16 | 805.74 | 60 | HOM | PASS | splice_acceptor_variant&intron_variant | DGKH |
| chr14 | 90062203 | G | GCCC | 30 | 6 | 29.74 | 60 | HET | MG_INDEL_Filter | start_lost&conservative_inframe_insertion | KCNK13 |
| chr15 | 23440078 | T | TG | 31 | 18 | 537.27 | 51.05 | HET | PASS | frameshift_variant | GOLGA6L2 |
| chr16 | 1241453 | G | A | 79 | 12 | 98.77 | 31.18 | HET | MG_SNP_Filter | structural_interaction_variant | TPSAB1 |
| chr16 | 30979994 | CTCCG | C | 87 | 9 | 43.73 | 60 | HET | MG_INDEL_Filter | frameshift_variant | SETD1A |
| chr17 | 69028583 | T | A | 118 | 11 | 12.99 | 60 | HET | MG_SNP_Filter | stop_gained | ABCA9 |
| chr19 | 1430254 | G | GGC | 21 | 4 | 76.73 | 60 | HET | PASS | frameshift_variant | DAZAP1 |
| chr19 | 8902218 | A | AG | 106 | 10 | 93.73 | 58.65 | HET | MG_INDEL_Filter | frameshift_variant | MUC16 |
| chr19 | 8902221 | AG | A | 104 | 10 | 87.73 | 58.62 | HET | MG_INDEL_Filter | frameshift_variant&splice_region_variant | MUC16 |
| chr19 | 35618400 | A | C | 61 | 8 | 156.77 | 60 | HET | PASS | splice_acceptor_variant&intron_variant | HAUS5 |
| chr19 | 35618401 | G | C | 62 | 6 | 132.77 | 60 | HET | PASS | splice_acceptor_variant&intron_variant | HAUS5 |
| chr19 | 39906057 | CCT | C | 12 | 12 | 592.73 | 42.03 | HOM | PASS | frameshift_variant | FCGBP |
| chr19 | 39906061 | A | AGG | 13 | 13 | 547.73 | 41.88 | HOM | PASS | frameshift_variant | FCGBP |
| chr19 | 39906132 | AGC | A | 24 | 24 | 1042.73 | 40 | HOM | PASS | frameshift_variant | FCGBP |
| chr19 | 39906137 | T | TGC | 24 | 24 | 1042.73 | 40 | HOM | PASS | frameshift_variant | FCGBP |
| chr19 | 39906156 | ACT | A | 17 | 17 | 862.73 | 40 | HOM | PASS | frameshift_variant | FCGBP |
| chr19 | 39906162 | C | CAGGGG | 20 | 20 | 907.73 | 44.49 | HOM | PASS | frameshift_variant | FCGBP |
| chr19 | 39906164 | A | AGG | 20 | 20 | 1042.73 | 45.6 | HOM | PASS | frameshift_variant | FCGBP |
| chr19 | 39906168 | GT | G | 22 | 22 | 1042.73 | 48.98 | HOM | PASS | frameshift_variant | FCGBP |
| chr19 | 41090053 | C | T | 170 | 19 | 134.77 | 58.48 | HET | MG_SNP_Filter | structural_interaction_variant | CYP2A13 |
| chr20 | 46128304 | C | CTT | 65 | 15 | 627.73 | 59.95 | HET | PASS | splice_acceptor_variant&intron_variant | CD40 |
| chr20 | 49850763 | G | GT | 158 | 17 | 39.73 | 60 | HET | MG_INDEL_Filter | frameshift_variant | SLC9A8 |
| chr22 | 37069345 | CTGGGG | C | 30 | 2 | 35.73 | 60 | HET | PASS | splice_acceptor_variant&splice_region_variant&intron_variant | TMPRSS6 |
| chr22 | 38245933 | A | C | 28 | 3 | 28.77 | 60 | HET | MG_SNP_Filter | splice_donor_variant&intron_variant | TMEM184B |
| chr22 | 39663712 | G | GCCCCC | 46 | 6 | 150.73 | 60 | HET | PASS | splice_acceptor_variant&intron_variant | CACNA1I |
| chrX | 1.01E+08 | GC | G | 19 | 9 | 210.73 | 58.83 | HET | PASS | frameshift_variant | ARMCX4 |
| chrX | 1.01E+08 | TGAGGC | T | 23 | 9 | 198.73 | 59.03 | HET | PASS | frameshift_variant | ARMCX4 |

**Supplementary Figures and Figure Legends**

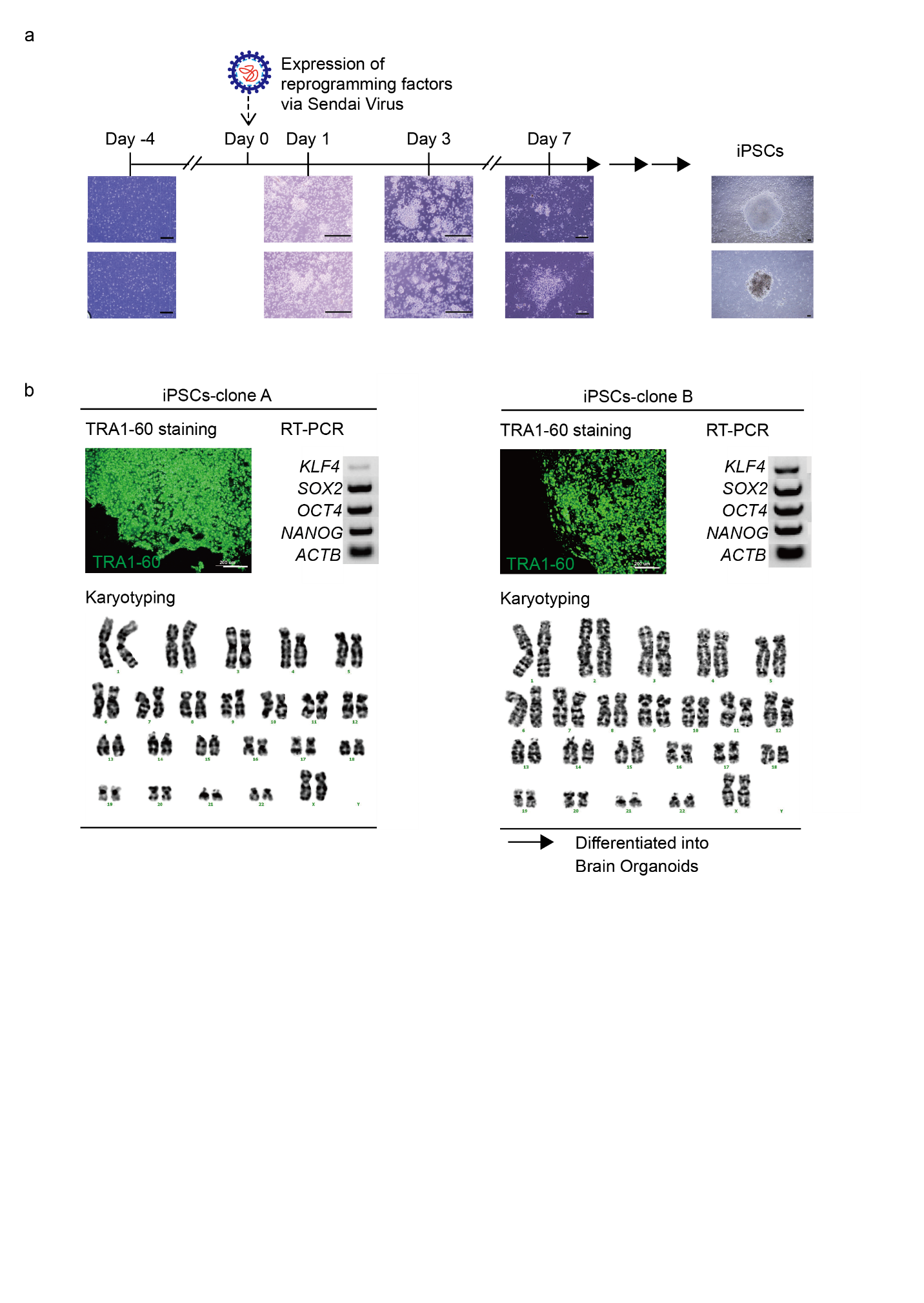

**Supplementary Figure 1. Generation and Characterization of iPSCs from PBMCs**

(A) Bright-field images showing the reprogramming process from PBMCs to the formation of iPSC colonies. Scale bar = 200μm.

(B) iPSC validation results. Live imaging of TRA1-60 staining of two iPSC colonies at passage 2. Scale bar = 200μm. Pluripotency marker expression (*KLF4*, *SOX2*, *OCT4*, and *NANOG*) was confirmed by RT-PCR, along with normal karyotype analysis, at passage 10. Among the two clones, clone B was used for organoid differentiation.

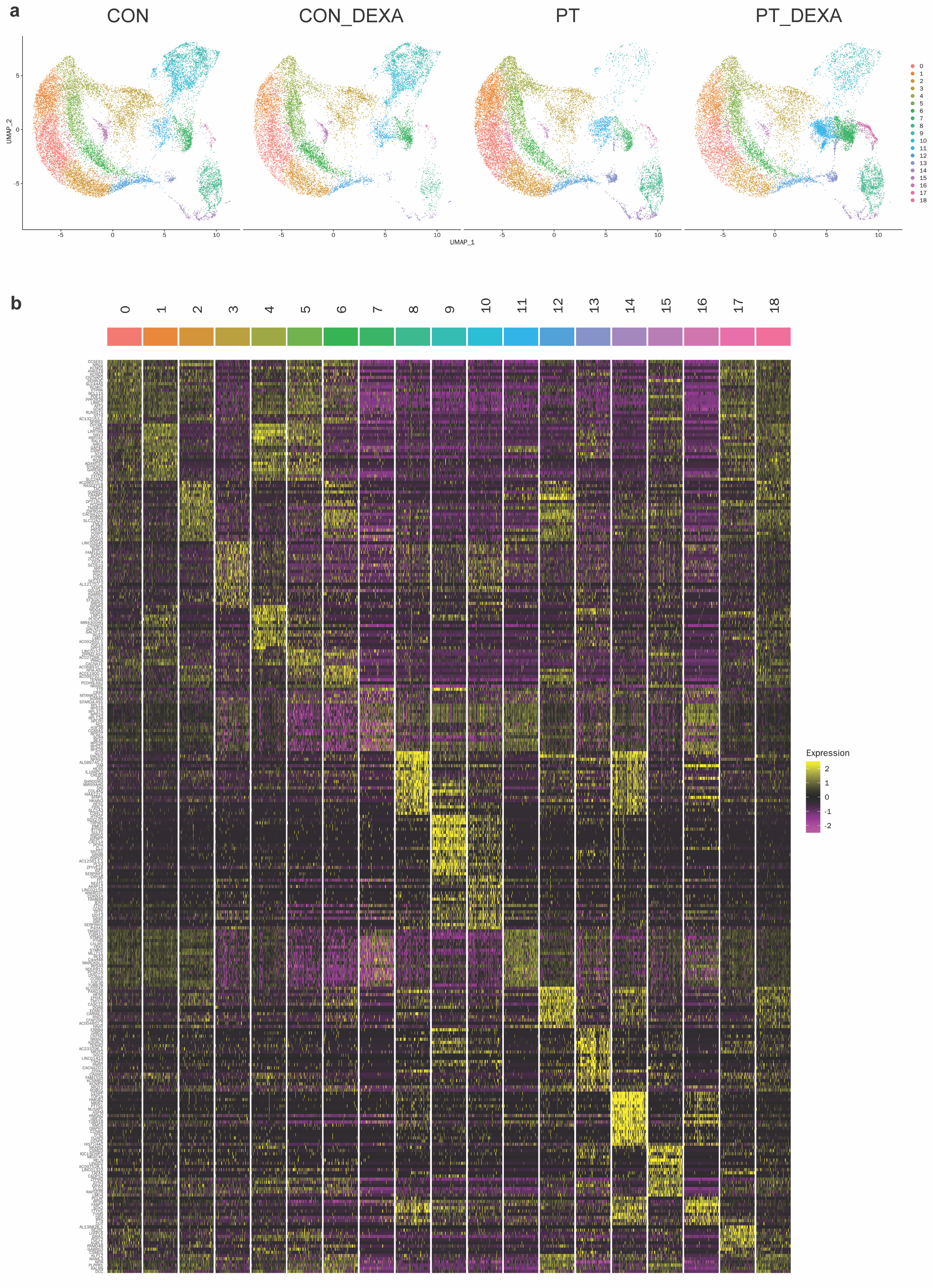

**Supplementary Figure 2. Single cell transcriptomic results of brain organoids of a patient and a control**

(A) Identified cell types through Seurat analysis from single-cell RNA sequencing of brain organoids and the distribution of cell types among brain organoids (n=3 for each group) from controls (CON) and patients (PT), both before and after dexamethasone (DMX) treatment.

(B) Heatmap showing the gene expression patterns for the top 20 genes in each cluster.

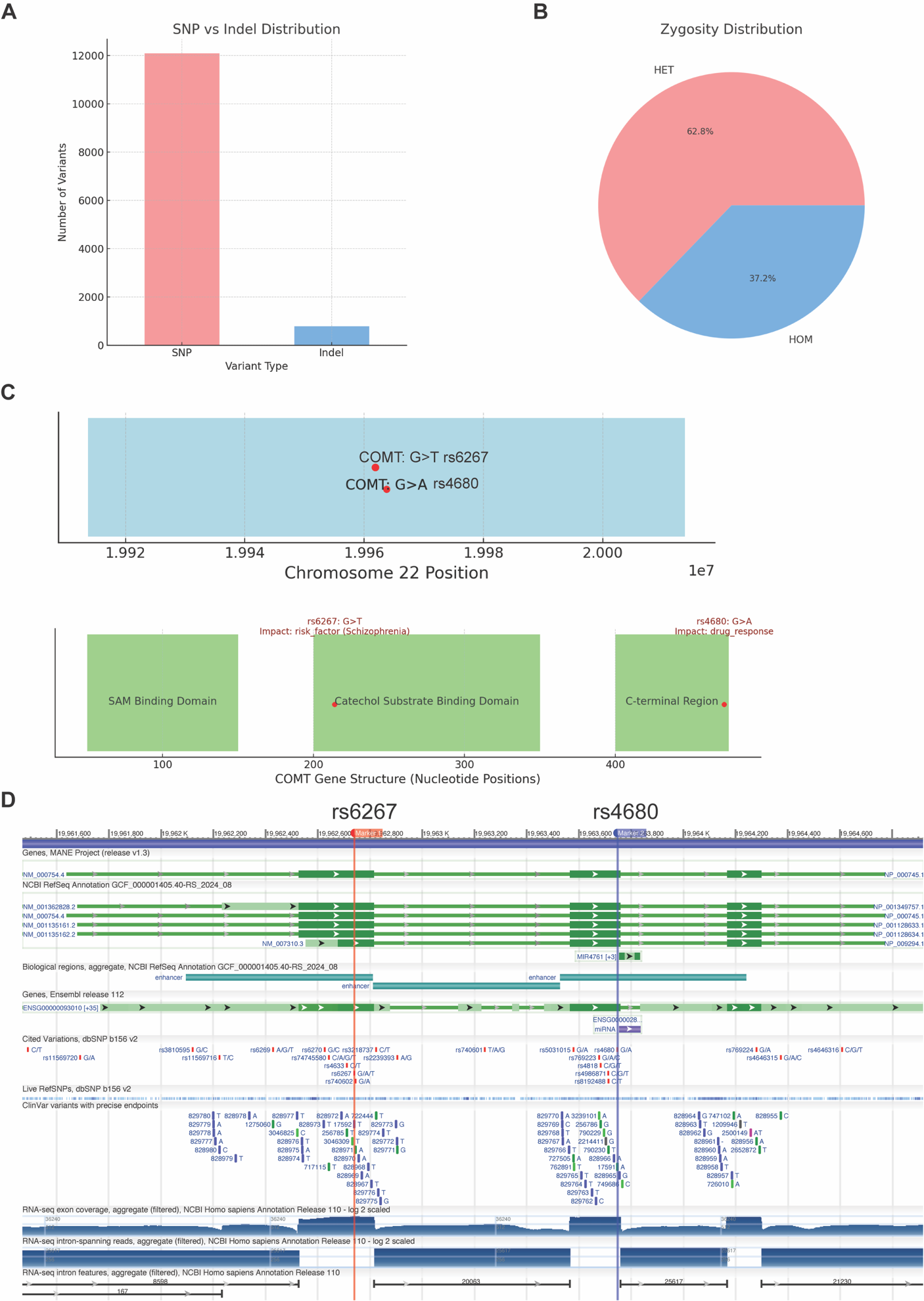

**Supplemenatry Figure 3. Whole-Exome Sequencing (WES) Reveals Genetic Polymorphisms Linked to Neurogenesis and Drug Response in a Patient with Atypical Depression and Psychotic Symptoms**

(A and B) Distribution of single-nucleotide polymorphisms (SNPs) and indels and zygosity distribution in WES results.

(C) Mutations on *COMT* at Chromosome 22 and mutations on *COMT* gene structures with domains.

(D) The detailed genomic organization of the *COMT* gene, including its transcript variants and associated SNPs such as rs4680 G>A and rs6267 C>T, based on NCBI RefSeq annotations.
